# Single cell transcriptomics in a treatment-segregated cohort exposes a STAT3-regulated therapeutic gap in idiopathic pulmonary fibrosis

**DOI:** 10.1101/2025.06.16.659944

**Authors:** Neil J. McKenna, Scott A. Ochsner, Alan Waich, Juan Cala-Garcia, Maria E. Echartrea, Sandra L. Grimm, Fernando Poli, Rafael C. Castillo, Juan D. Zuluaga, Sergio Poli, Taylor S. Adams, Ricardo Pineda, Benjamin Moss, Tianshi D. Wu, Charles G. Minard, Fadi Nikola, Laura Fernandez, Stefan W. Ryter, Rudolf T. Pillich, Julian A. Villalba, Kosuke Kato, Louise Hecker, Lindsay J. Celada, Maor Sauler, Gabriel Loor, Melanie Konigshoff, Naftali Kaminski, Benjamin Raby, Sandeep K. Agarwal, Konstantin Tsoyi, Cristian Coarfa, Ivan O. Rosas

**Author notes:** Corresponding author: Ivan O. Rosas, MD, Section of Pulmonary, Critical Care and Sleep Medicine Department of Medicine, One Baylor Plaza, Baylor College of Medicine Houston, TX 77030, USA, e. Yale School of Medicine, New Haven, CT, USA. Department of Medicine, University of Maryland Baltimore, Baltimore, MD, USA. One Sentence Summary: STAT3 is a prominent and targetable driver of clinical IPF transcriptional programs that are refractory to approved first generation antifibrotics.

## Abstract

Idiopathic pulmonary fibrosis (IPF) is a progressive fibrotic pulmonary disease of unknown etiology. Since approved IPF drugs only slow disease progression, novel therapeutics are required that improve clinical outcomes. Here we report a single cell lung RNA-Seq and gene regulatory network analysis of the largest IPF cohort assembled to date. Segregating this cohort based on status of treatment with approved first-generation IPF antifibrotics (untreated, nintedanib-and pirfenidone-treated), we describe for the first time the transcriptional landscape of untreated IPF across 40 lung cell types, and the elements of this program that are impacted by these antifibrotics. On average, nearly 60% of the untreated IPF-dysregulated transcriptome is refractory to treatment with these drugs, a transcriptional deficit we refer to as the IPF therapeutic gap. Gene regulatory network analysis indicated a dominant functional footprint for the transcription factor STAT3 in both untreated IPF and the IPF therapeutic gap. Validating our analysis in a translational precision cut lung slice platform that recapitulates IPF explants, pharmacological inhibition of STAT3 reduced the IPF therapeutic gap in numerous lung cell types. Finally, we resolved a STAT3-anchored master regulatory network comprising numerous profibrotic transcription factors in IPF alveolar fibroblasts, a critical fibrotic lineage. Our study represents a comprehensive resource for translational lung fibrosis research and introduces a strategy for drug discovery that is adaptable to human disease more broadly.

## INTRODUCTION

Idiopathic pulmonary fibrosis (IPF) is a chronic lung disease characterized by progressive decline of lung function leading to respiratory failure (*1*). The underlying mechanisms of IPF pathogenesis remain unclear but may be initiated by alveolar epithelial cell injury leading to immune, stromal and endothelial cell recruitment, fibroblast activation and dysregulated extracellular matrix (ECM) production that ultimately remodels lung architecture (*2*). Two small molecules have been approved for treatment of IPF, nintedanib and pirfenidone. Nintedanib is a broad-spectrum receptor tyrosine kinase inhibitor originally developed as a non-small cell lung cancer therapeutic (*3*), whereas pirfenidone has been shown to reduce the activity of growth factor and p38 MAPK-regulated pathways (*4*). Although their direct protein targets have been characterized to some extent, the transcriptional events affected by these first-generation drugs in specific IPF cell types remain largely undefined at scale. Transcriptional insights into the mechanism of action of nintedanib and pirfenidone derived from cell culture and animal models have limited direct relevance to clinical end points. The consequent lack of information on specific transcriptional programs impacted by these drugs in patient lungs has hampered the development of novel therapeutics in IPF. We and others have previously published single cell-resolution studies that have documented transcriptional programs underlying the etiology of IPF (*5–8*). Since these IPF cohorts comprised subjects who had undergone therapy using approved antifibrotics, they were not designed to discriminate between gene expression programs in untreated and treated IPF, nor to illuminate the transcriptional impact of IPF therapies. Here we describe a data-driven approach to targeting transcription factors (TFs) that are strongly implicated in the development of clinical IPF and that are refractory to treatment with approved antifibrotics.

## RESULTS

### A treatment status-based single cell transcriptional analysis of IPF

We established the largest IPF treatment cohort to date, then segregated it according to treatment status prior to transplant surgery. This cohort consisted of lungs from 22 untreated, 24 pirfenidone-treated and 28 nintedanib-treated IPF patients, and 64 control lungs (**Fig. 1A**). This stratification enabled us to (i) define untreated IPF-dysregulated transcriptional programs; (ii) determine their regulation by approved antifibrotics; and (iii) identify IPF biological networks that were refractory to approved antifibrotic treatments. **Table S1** describes the subject characteristics for our study. We defined 40 explant Human Lung Cell Atlas cell types based on distinct gene markers (**fig. S1**). For brevity we assigned acronyms to the 40 cell types (**table S2**). To maximize statistical power for detecting differential gene expression, we combined our new cohort with that from our previous IPF single cell RNA sequencing (scRNA-Seq) analysis

**Figure 1.**
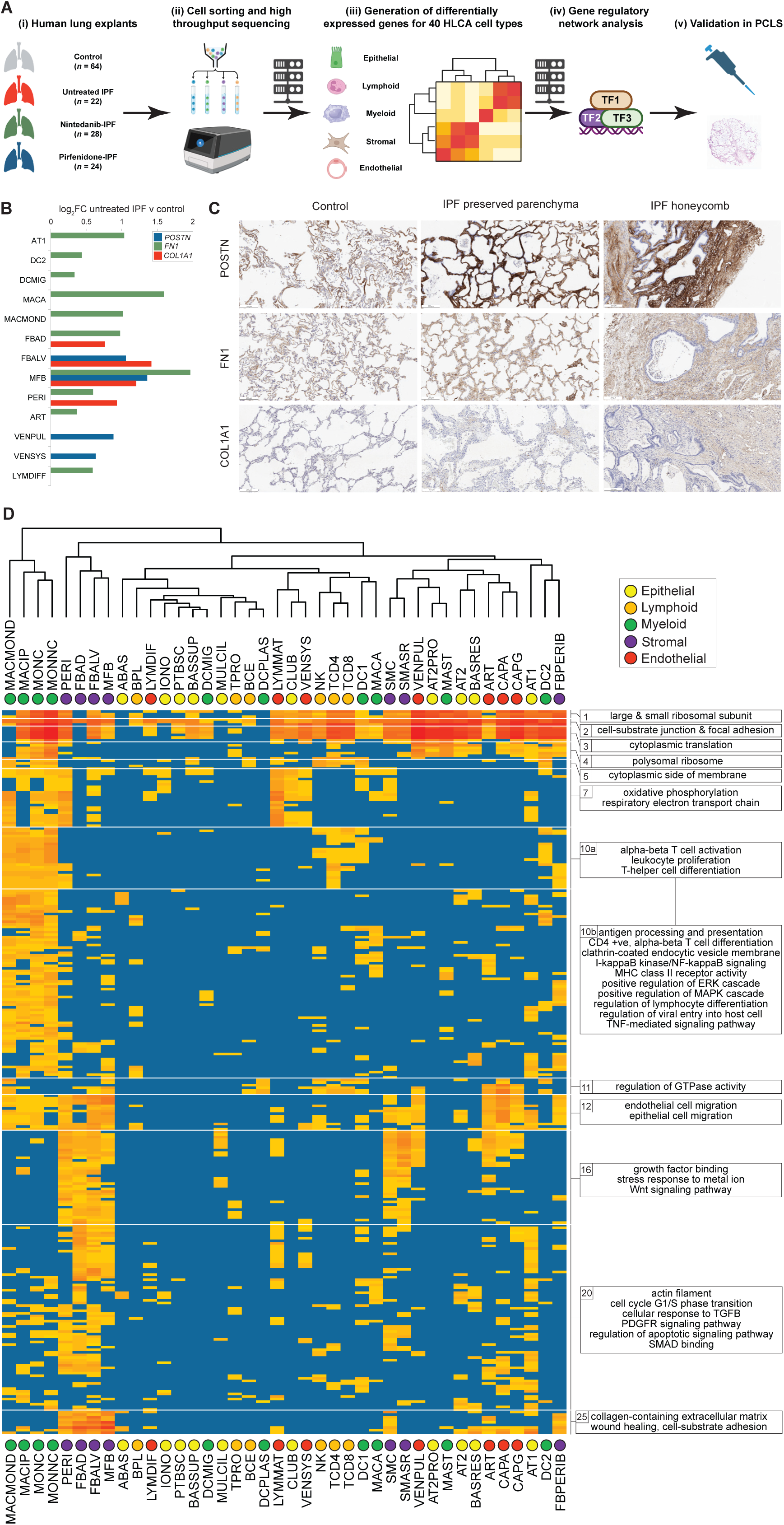
A treatment status-based single cell transcriptional analysis of IPF. Refer to **table S2** for cell type acronym definitions. **(A)** Overview of experimental design. **(B)** Horizontal bar chart showing untreated IPF v control induction of *POSTN*, *FN1* and *COL1A1* by cell type. **(C)** Representative immunohistochemical images of normal parenchyma (left), IPF preserved parenchyma (middle) and IPF honeycomb (right) stained for POSTN (upper), FN1 (middle) and COL1A1 (right). Scale bar represents 200µm. **(D)** Hierarchical clustering of selected enriched GO terms in untreated IPF-induced gene sets. Refer to **table S2** for cell type acronym definitions. GO enrichment analysis was performed on genes induced in untreated IPF (log_2_FC>0.32, FDR<0.1) relative to controls. Color palette represents-log10p enrichment scale from *p* adj < 0.05 (light orange) to *p* adj <1E-70 (red); blue indicates no significant enrichment. Terms that were significantly enriched (*p* adj < 0.05) in at least six cell types were subjected to hierarchical clustering as described in the methods section. Full heatmap is in **fig. S6** and full tabular data are in **table S6**.

(*5*). Batch effects between the two datasets were addressed using BBKNN (*9*) and scVI (*10*) and both methods were evaluated using scib-metrics (*11*). As shown in **fig. S2** and **table S3**, BBKNN and scVI demonstrated comparable performance across a broad set of integration metrics, supporting the robustness of our downstream analyses.

### The transcriptional landscape of untreated IPF at single cell resolution

As a preliminary step we used scCoda (*12*) to assess the effect of approved antifibrotics on explant cell type counts (**fig. S3** and **table S**4). Excepting an increase in the number of multiciliate cells in response to nintedanib treatment, no significant effect of antifibrotics on cell counts was observed. We next generated differentially-expressed genes (DEGs; log_2_ FC>0.32 or <-0.32, FDR<0.1) for three treatment-segregated subcohort contrasts (untreated IPF vs. control, pirfenidone-treated IPF vs. untreated IPF and nintedanib-treated IPF vs. untreated IPF) across the 40 explant cell types. The explant scRNA-Seq analysis was organized into a 1,494,240 data point DEG matrix documenting relative abundance values for 12,452 genes across the 40 cell types in the three subcohort contrasts (**table S5**). Genes in **table S5** are annotated using the Signaling Pathways Project (SPP) knowledgebase classification (*13*), which provides for filtering of genes according to signaling node category (e.g. receptors, enzymes, transcription factors) class (e.g. ligands, kinases, BZIP factors) and family. We observed no appreciable effect of covariates (age, sex, smoking status) on explant DEGs for all three contrasts across most cell types (**fig. S4**). Verifying broad alignment of our analysis with existing independent single cell transcriptomic studies of IPF, we observed robust overlap between untreated IPF-induced genes from selected clusters in our own analysis and IPF-induced signatures of comparable cell types from the Reyfman (*8*) and Haberman (*6*) IPF cohorts (**fig. S5**). We generated a pan-lung untreated IPF inductome that ranks genes by number of cell types with significant untreated IPF v control induction and average log_2_FC (**table S5**, columns AV-AX). Aligning our analysis with existing knowledge in the IPF field, the inductome assigned prominent rankings to numerous genes with established connections to lung fibrosis, including *POSTN, FN1, COL1A1* and others. As anticipated, particularly strong induction of these genes was observed in stromal cell types, with additional peaks in cells of the endothelial compartment (**Fig. 1B**). As an additional validation step, immunohistochemical analysis of POSTN, FN1 and COL1A1 verified strong induction of these markers at the protein level in IPF tissue relative to controls (**Fig. 1C** and **fig. S6A**).

To resolve cellular pathways dysregulated in untreated IPF, we performed Panther GO enrichment analysis on genes induced in untreated IPF vs. control (log_2_FC>0.32, FDR< 0.1) across the 40 cell types (**table S6**). We then conducted K-means clustering of untreated IPF-induced GO terms that were enriched (*p* adj < 0.05) in at least six cell types (*n* = 508), then partitioned the analysis at k=25 to resolve a set of 25 GO term clades. **Figure 1D** shows selected clades from this analysis; the full heat map is in **fig. S6B.** Clades 1-5 reflected strong induction across multiple compartments of genes with roles in ribosomal biogenesis, which contributes to signature IPF processes such as epithelial to mesenchymal transition, fibroblast to myofibroblast differentiation and T cell activation (*1*). Reflecting the established connection between cellular metabolism and fibrosis, mitochondrial oxidative phosphorylation and respirasome terms (clade 7) were present in endothelial and epithelial cell types, as well as in the myeloid and stromal compartments. The strong immune cell character of terms in clades 10a and 10b was consistent with their enrichment peaks in lymphoid and myeloid cell types (**Fig. 1D**). Consistent with its role in endothelial cell migration and dysfunction in pulmonary fibrosis (*14*), we observed peaks of GTPase signaling (clade 11) and endothelial cell migration (clade 12) enrichment in cells of the endothelial compartment. Clades that were largely restricted to the stromal compartment encompassed processes with strong connections to IPF, including growth factor and Wnt signaling (clade 16); TGFB, PDGFR and SMAD signaling in clade 20; and collagen-containing ECM and wound healing in clade 25 (**Fig. 1D**) (*1*).

### Global impact of approved antifibrotics in IPF

We next evaluated the effects of nintedanib and pirfenidone on the untreated IPF transcriptome. We computed the percentage of untreated IPF-DEGs (induced and repressed) that were reversed by antifibrotics in each cell type (cutoffs: log_2_FC >0.32 or <-0.32, FDR < 0.1).

Remarkably, we found that on average, nearly 60% of untreated IPF DEGs were not reversed by treatment with either antifibrotic **(Fig. 2A; table S7**). Because it represents a gap between desired and actual performance of approved therapeutics, we refer to this transcriptional deficit as the IPF therapeutic gap. The size of the gap varied from >95% in ionocytes and plasmacytoid dendritic cells to less than 25% in CD4^+^ T and natural killer cells (**Fig. 2A** and **table S7**). Averaged across cell types, the gap was largest in the epithelial compartment (65%), followed by myeloid (62%), stromal (57%) and endothelial (53%), with the smallest gap observed in the lymphoid compartment (52%; **table S7**). Although the average overlap between nintedanib-and pirfenidone-regulated genes was modest (∼20% on average), in certain cell types it was much higher (TCD4, 51%; LYMMAT, 43%, TCD8, 40%; **table S7**), indicating significant redundancy of function of approved antifibrotics in these cell types.

**Figure 2.**
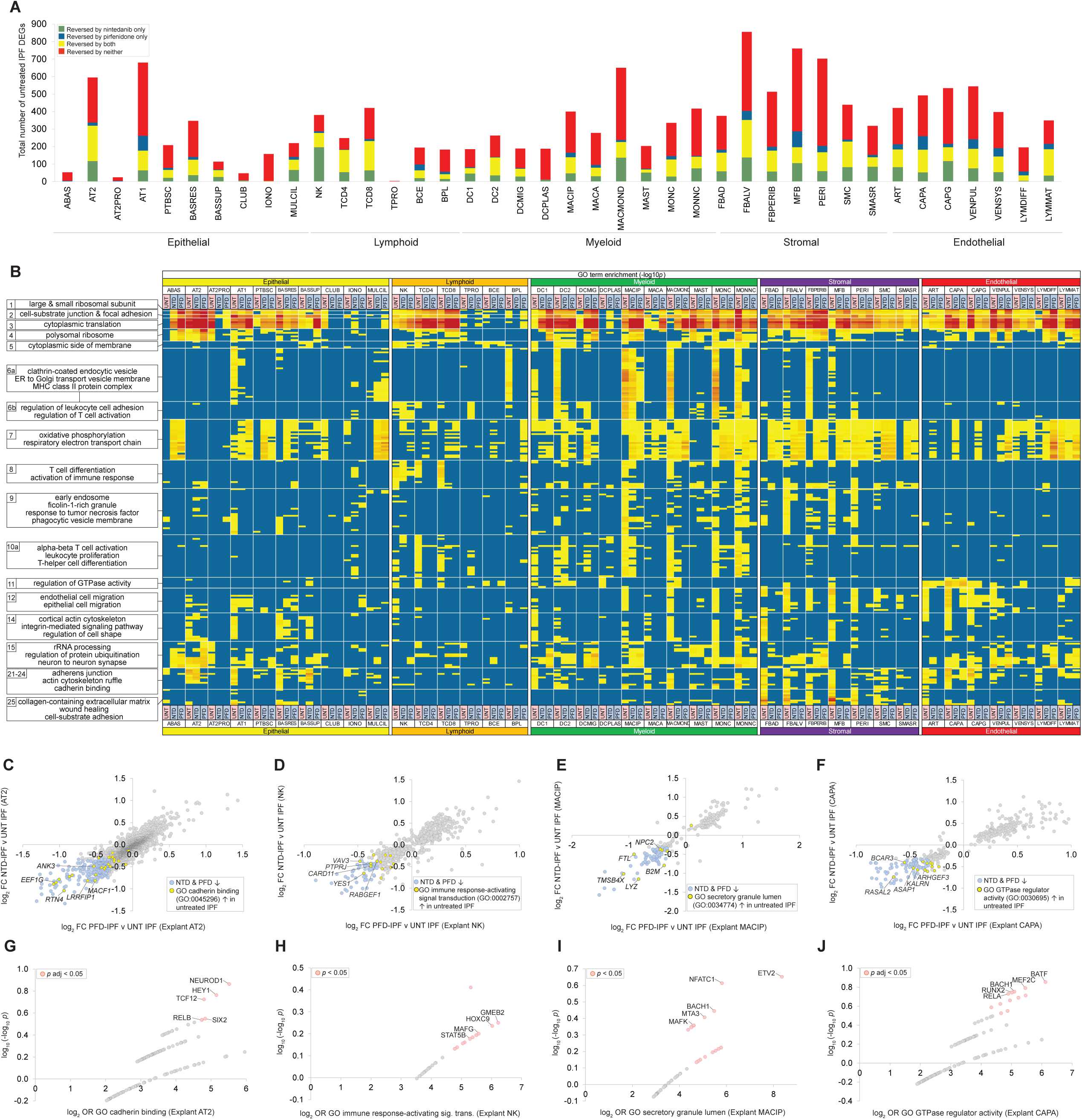
Single cell transcriptional biology of approved antifibrotics in IPF. Refer to **table S2** for cell type acronym definitions. **(A)** Stacked bar chart representation of differentially expressed genes derived from explant scRNA-Seq data. Bars represent total numbers of untreated IPF vs. control-dysregulated (log_2_ FC>0.32 or <-0.32, FDR<0.1) genes in each cell type at the indicated cutoffs. Subsets of these genes whose dysregulation is reversed by nintedanib only (green), pirfenidone only (blue), both antifibrotics (yellow) and neither (red) are indicated. Full data are in **table S7**. **(B)** Heatmap indicating the extent to which untreated IPF-induced GO terms are reversed by treatment with nintedanib or pirfenidone in each cell type. Selected GO term clades from **Fig. 2** are arranged on the vertical axis; the 40 explant lung cell types are arranged into their five compartments of origin on the horizontal axis. On the horizontal axis label, for a given cell type-GO term clade intersection, UNT (red) indicates enrichment of that GO term in untreated IPF-induced genes; NTD (blue) indicates enrichment of that GO term in nintedanib-IPF-repressed genes; and PFD (blue) indicates enrichment of that GO term in pirfenidone-IPF-repressed genes. Within the heatmap, the color palette represents - log10*p* enrichment scale from *p* adj < 0.05 (yellow) to *p* adj <1E-25 (red); blue indicates no significant enrichment. Full tabular data are in **table S6**. Panels **C-F** represent scatterplots of pirfenidone-IPF v untreated IPF (*x* axis) and nintedanib-IPF vs. untreated IPF (*y* axis) log_2_ FC values for untreated IPF-induced genes mapped to the following GO terms: **(C)** cadherin binding in AT2 cells; **(D)** immune response-activating signal transduction in NK cells; **(E)** secretory granule lumen in MACIPs; and **(F)** GTPase regulator activity in CAPA cells. Panels **G-J** represent GRN plots generated by HCT intersection analysis of gene sets highlighted in blue in panels **C-F**.

### Single cell transcriptional biology of approved antifibrotics in IPF

To investigate the biology of approved antifibrotics in IPF, we evaluated the extent to which dysregulation of functional ontologies in untreated IPF was reversed by antifibrotic treatment. As in **Fig. 2**, we focus here on untreated IPF-induced functional ontologies (see **table S6** for full data). We performed Panther GO enrichment analysis on (i) nintedanib-treated IPF vs. untreated IPF-repressed gene sets; and (ii) pirfenidone-treated IPF vs. untreated IPF-repressed gene sets (log_2_FC <-0.32, FDR < 0.1) and compared the results with the GO analysis of untreated IPF-induced gene sets. **Figure 2B** shows enrichment of selected GO terms in (i) untreated IPF-induced, (ii) nintedanib-repressed and (iii) pirfenidone-repressed gene sets. Our analysis indicated robust effects of antifibrotics across all compartments on ribosomal biogenesis, cell-substrate junction and focal adhesion (clades 1-5) and OxPhos (clade 7). The latter is consistent with reported effects of approved antifibrotics on oxidative stress (*15*).

Similarly, repression by approved antifibrotics of terms related to regulation of GTPase activity (clade 11) as well as epithelial and endothelial cell migration (clade 12) was evident. In general, and reflecting the DEG breakdown by therapy status (**Fig. 2A**), nintedanib exerted a stronger effect on untreated IPF-induced programs compared to pirfenidone.

Repressive effects of approved antifibrotics on specific untreated IPF-induced GO term-mapped genes in selected cell types are shown in **Fig. 2C-F**. Strong effects of antifibrotics were observed on cellular processes previously implicated in IPF progression, including cadherin binding in AT2 cells (clade 22; **Fig. 2C**), immune signal transduction in natural killer cells (clade 8; **Fig. 2D**), secretory granule processing in interstitial macrophages (clade 9; **Fig. 2E**), and Ras GTPase signaling in capillary aerocyte endothelial cells (clade 11; **Fig. 2F**). In several instances, antifibrotics impacted untreated IPF-dysregulated processes in a cell type-specific manner. For example, Golgi and MHC II-related terms (clade 6a) were at least partially repressed by both antifibrotics in MACIP cells (**Fig. 2B&3E**) but not in other myeloid cell types.

Gene regulatory networks (GRNs) are contextual sets of transcriptional regulators that drive specific gene expression programs in response to afferent signaling events (*16*). We previously described a curated public TF ChIP-Seq dataset-based approach to predicting GRNs, referred to as high confidence transcriptional target (HCT) intersection analysis (**fig. S7**)(*17*). To predict GRNs targeted by approved antifibrotics to effect repression of the GO term-mapped genes in **Fig. 2C-F**, we carried out HCT intersection analysis on those mapped genes that were induced in untreated IPF and repressed by both antifibrotics in the indicated cell type. The GRN composition for GO cadherin binding genes in AT2 cells reflected previously characterized roles of Notch/HEY1, TCF12, SIX2 and RELB/p65 in regulation of cadherin biology **(Fig. 2G)**.

Similarly, previous studies have implicated HOXB13, HOXC9, MAFG and STAT5B in NK cell activation **(Fig. 2H)**, and ETV2, NFATC1, BACH1, MTA3 and MAFK in macrophage activation and differentiation **(Fig. 2I)**. Finally, elevated rankings assigned to BATF, MEF2C, GATA4, BACH1 and RELA in the CAPA analysis align with existing studies implicating these TFs in endothelial cell morphology and function **(Fig. 2J)**. In addition to indicating broad overlap between nintedanib and pirfenidone-regulated genes in numerous cell types, our analysis also exposed pathways and regulatory networks that were selectively impacted by either drug (**fig. S8**).

### STAT3 dominates untreated IPF-induced gene regulatory networks

We next generated a pan-lung GRN to identify TFs that had the strongest functional footprints across all untreated IPF cell types. To achieve this, we performed HCT intersection analysis on individual untreated IPF-induced gene sets from the 40 cell types. TFs were scored by number of cell types containing a significant footprint and mean footprint-log_10_*p*, then ranked on this score. The full data are shown in **table S8**, again annotated using the SPP vocabulary (*13*). **Fig. 3A** shows the top 25 TFs prioritized by this analysis. Validating our analysis, a set of TFs with established roles in pulmonary fibrosis curated by the FibROAD resource (*18*) was strongly enriched within the top ranked TFs in the pan-lung GRN (**Fig. 3B** and **table S8,** column AW). In addition to indicating a functional hierarchy for TFs across the IPF lung, our analysis affords insight into the diversity of composition of IPF-induced GRNs in distinct lung cell types. The AT2 GRN contained strong footprints for pioneer TFs with established roles in lung development and morphology, such as NKX2-1 and members of the FOXA and GATA families, in addition to STAT3 (**fig. S9A** and **table S8**, column G). Elevated rankings in the CD4+ T cell GRN were assigned to TFs with key roles in lymphoid lineage biology, including FOXP1, RELB, NFATC1, BCL6 and MTA3 (**fig. S9B** and **table S8**, column Q). In addition, functional footprints for TFs connected to endothelial cell lineages, such as NFATC1, NFIC, MEF2A, TEAD1 and TEAD4 dominated the capillary aerocyte endothelial GRN (**fig. S9C** and **table S8**, column AN).

**Figure 3.**
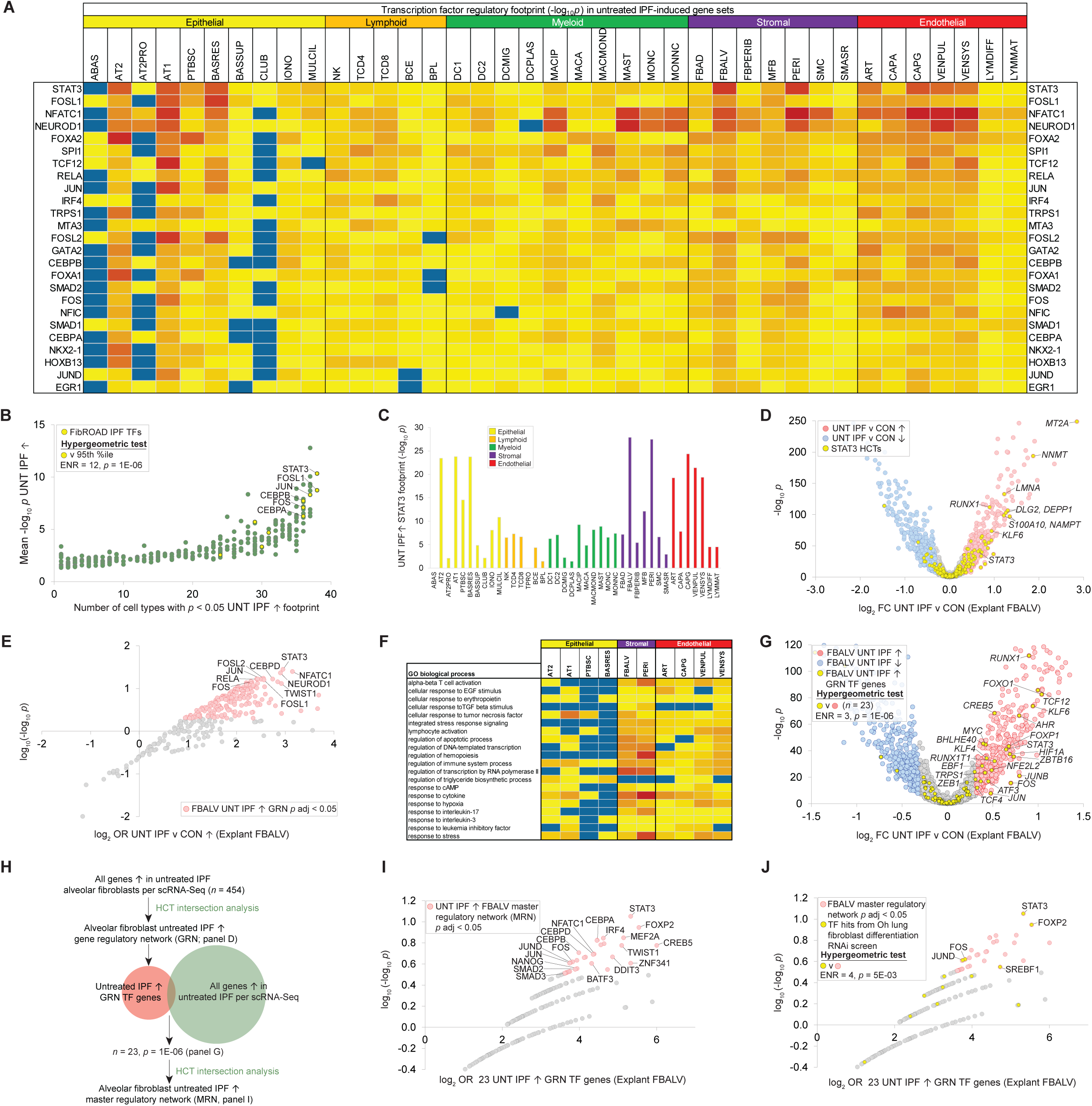
STAT3 is a dominant transcription factor in untreated IPF. Refer to **table S2** for cell type acronym definitions. **(A)** Heat map showing predicted top 25 TFs in untreated IPF-induced pan-lung GRN. HCT intersection analysis was performed on untreated IPF-induced genes in the 40 cell types. TFs are ranked based on (i) number of significant footprints and (ii) average footprint size (-log_10_*p*). Full data are in **table S8**. Color palette represents-log_10_*p* enrichment scale from *p* adj < 0.05 (yellow) to *p* adj <1E-35 (red); blue indicates no significant enrichment. TPRO cells had no top 25 TF footprints and are omitted. **(B)** Validation of pan-lung GRN using a panel of FibROAD-(*18*) curated TFs with established roles in pulmonary fibrosis (**table S8**, column AW). *x* axis: number of significant footprints; *y* axis: average footprint size. **(C)** Predicted STAT3 HCT footprints in untreated IPF-induced genes across the 40 cell types. **(D)** Volcano plot showing distribution of predicted STAT3 HCTs in untreated IPF vs. control DEGs in alveolar fibroblasts. **(E)** Untreated IPF-induced alveolar fibroblast GRN. TFs with the strongest and most significant footprints within the gene set of interest are distributed towards the top right of the plot. **(F)** Heat map showing selected GO biological processes enriched among untreated IPF-induced STAT3 HCTs in the top ten cell types ranked by STAT3 footprint. Full data are in **table S10**. Color palette represents-log10*p* enrichment scale from *p* < 0.05 (yellow) to *p* <1E-06 (red); blue indicates no significant enrichment. **(G)** Volcano plot of untreated IPF vs. control explant alveolar fibroblast DEGs highlighting genes encoding IPF-induced GRN TFs. **(H)** Informatic strategy for generating explant alveolar fibroblast MRN. **(I)** Untreated IPF-induced explant alveolar fibroblast MRN. Full data are in **table S11**. **(J)** Untreated IPF-induced explant alveolar MRN highlighting TFs encoded by genes that were hits in an RNAi screen of lung fibroblast differentiation (7).

The TF with the broadest predicted transcriptional footprint in untreated IPF-induced genes across all cell types was STAT3 (**Fig. 3A**), which we previously demonstrated was aberrantly activated in experimental and clinical pulmonary fibrosis (*19*). Corroborating our findings, we observed strong identity between the top ranked TFs in the pan-lung GRN and those in a GRN analysis of a published dataset (GSE32537) of genes induced in severe v mild IPF (**fig. S9D**), with STAT3 ranked first in both analyses (**table S9**). The strongest predicted STAT3 untreated IPF-induced functional footprint across the 40 cell types was in alveolar fibroblasts (**Fig. 3C**), a cell type implicated as the primary source of fibroblast subtypes in response to lung injury (*20*). Other peaks of predicted STAT3 activity were observed in cell types in the epithelial (AT2, AT1, resting basal epithelial cells) and endothelial (artery, general capillary and venous systemic) compartments (**Fig. 3C**). The predicted STAT3 footprint in alveolar fibroblast untreated IPF-induced genes (**Fig. 3D**) included numerous genes implicated in fibrosis in the lung and other organs, including *MT2A*, *NNMT*, *LMNA*, *NAMPT* and *STAT3* itself. In addition to STAT3, elevated rankings in the alveolar fibroblast untreated IPF-induced GRN were assigned to other known profibrotic TFs, including NFATC1, TWIST1 and members of the CEBP and AP-1 TF families **(Fig. 3E** and **table S8).** Of note, we observed very strong identity (*r*^2^ = 0.73) in the composition of the alveolar fibroblast untreated IPF-induced GRN and that in pericytes (**fig. S9E**), another stromal cell type known to contribute to IPF pathogenesis (*21*).

To investigate STAT3-regulated biological pathways across the lung, we carried out Panther GO analysis on untreated IPF-induced STAT3 HCTs (annotated in **table S5,** column G) in the top ten explant cell types ranked by size of STAT3 footprint. **Figure 3F** shows selected pathway enrichments from this analysis; the full data are in **table S10**. Ontologies predicted to be regulated by STAT3 were functionally diverse and encompassed numerous pathways with established roles in pulmonary fibrosis, including the stress response, apoptosis and hypoxia (**Fig. 3F**). The established role of STAT3 as a transcriptional mediator of a broad spectrum of cellular signaling inputs was reflected in enrichment of pathways regulated by interleukins (IL-3, IL-17), cytokines (TNF), growth factors (EGF, TGFβ, LIF) and cAMP. Of note, although no immune cells were among the top ten cell types, immune-related ontologies featured prominently in our analysis, potentially indicating STAT3-mediated crosstalk between the epithelial, stromal and vascular endothelial cell types and the immune compartment (**Fig. 3F**).

### STAT3 anchors a master regulatory network in IPF alveolar fibroblasts

As the cell type with the largest IPF-induced STAT3 footprint, we focused our attention on alveolar fibroblasts. Because TFs gain function in part through increased abundance, we hypothesized that gain-of-function of TFs in IPF alveolar fibroblasts encompassed transcriptional induction of their encoding genes. Consistent with our hypothesis, genes encoding 23 members of the IPF-induced GRN were significantly enriched among genes transcriptionally induced in untreated IPF alveolar fibroblasts (ENR = 3, *p* = 1E-06; **Fig. 3G; table S5**, column AY and **table S8**, column AX). GRN footprints for the 23 TFs encoded by these genes are shown in **fig. S9F**. Notably, the set of 23 TF genes was strongly enriched (ENR = 72, *p* = 1E-10) for a set of experimentally-defined immediate early response genes (*22*), considered a hallmark of growth factor stimulation (**fig. S9G**). Master regulatory networks (MRNs) supervise GRNs in specific cell types and states by modulating both expression and function of GRN TFs (*23, 24*). To characterize the MRN controlling the expression of genes encoding GRN TFs in IPF alveolar fibroblasts, we performed HCT intersection analysis on the set of 23 GRN TF genes that were transcriptionally induced in untreated IPF in this cell type (**Fig. 3H**). STAT3 had the strongest footprint in the MRN (**Fig. 3I** and **table S11**). Surveying the MRN, we identified known STAT3-interacting TFs, including NANOG, AP-1 family members, DDIT3, MEF2C and TWIST1. Moreover, of the *p* adj < 0.05 GRN members, BATF3, IRF4 and STAT3 are in a core regulatory network previously identified in anaplastic large cell lymphoma (*25*) (ENR = 15, *p* = 1E-04; hypergeometric test). In addition, a set of TF hits from an RNAi screen of lung fibroblast to myofibroblast differentiation (*26*), which included STAT3, FOS and JUND, was enriched within the MRN (ENR = 5, *p* = 5E-03; **Fig. 3J**). Furthermore, the MRN was appreciably enriched for orthologs of TFs encoded by genes whose null deletion results in defective wound healing in mice (**fig. S9H**). In summary, we identified STAT3 as a dominant TF across the untreated IPF lung, with a particularly influential regulatory role in alveolar fibroblasts, a critical pathological IPF lineage.

### Characterization of the IPF-induced therapeutic gap

Since it represented a logical target for the development of novel antifibrotics, we next wished to characterize the biology of the IPF therapeutic gap. Due to space considerations, we focus here on the IPF-induced therapeutic gap. To characterize this gap, for each cell type we identified genes induced in untreated IPF but not repressed by either approved antifibrotic (**table S12**).

We then performed GO analysis on these genes and ranked terms based on (i) the number of significant cell type enrichments and (ii) mean enrichment-log_10_*p* across the 40 cell types (**table S13**, columns AS-AU). We repeated this ranking step on the GO terms generated by our previous analysis of all untreated IPF-induced genes (**table S6**, columns AU-AW); a total of 4439 terms were common to both analyses. To prioritize terms with a higher ranking in the IPF-induced gap analysis than in the analysis of all untreated IPF-induced genes, we next calculated the ratio (untreated IPF-induced GO rank/IPF-induced gap GO rank; **table S13**, column AW) and ranked terms on this value. **Figure 4A** shows the top 30 ranked gap-prioritized terms.

**Figure 4.**
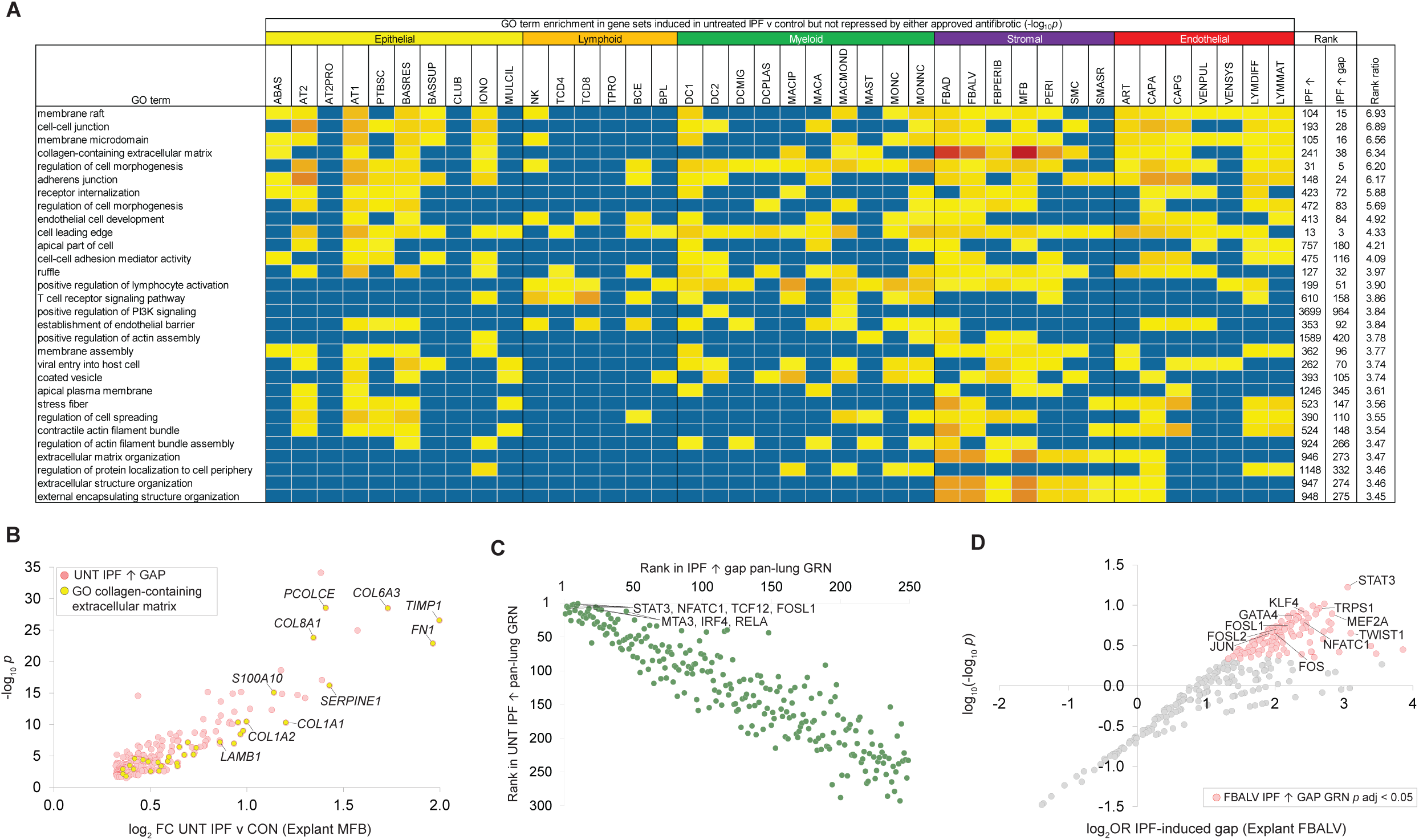
Characterization of the IPF-induced therapeutic gap. Refer to **table S2** for cell type acronym definitions. **(A)** Heatmap showing enrichment of top 30 prioritized GO terms in the IPF-induced therapeutic gap (i.e. genes induced in untreated IPF but not repressed by either antifibrotic) in each cell type. GO terms were prioritized as described in the text. Color palette represents-log10*p* enrichment scale from *p* adj < 0.05 (yellow) to *p* adj <1E-25 (red); blue indicates no significant enrichment. Full data are in **table S13**. **(B)** Induction in untreated IPF vs. control explant myofibroblasts of IPF-induced gap genes mapping to GO collagen-containing ECM. Full data are in **table S12**. **(C)** Scatterplot comparing TF rankings in the untreated IPF-induced (*x* axis) and IPF-induced gap (*y* axis) pan-lung explant GRNs. Full gap GRN data are in **table S14**. **(D)** GRN plot for the untreated IPF-induced gap in explant IPF alveolar fibroblasts.

Validating our approach, the GO terms that we had identified as strongly regulated by approved antifibrotics, such as mitochondrial OxPhos and ribosomal biogenesis (**Fig. 2B**), were among the lowest ranked terms. The top gap-prioritized terms emerging from this analysis were strongly represented in stromal, endothelial and epithelial cell types (**Fig. 4A**). Consistent with the comparatively strong effects of approved antifibrotics observed in the lymphoid and myeloid compartments (**Fig. 2B**), enrichment of GO terms in the IPF-induced gap in these cell types was relatively low. Prioritization in the IPF-induced gap of the term “collagen-containing extracellular matrix” (ranked 4/4439 GO terms) was surprising, indicating as it did that approved antifibrotics only modestly impacted IPF induction of these genes. As shown in **Figure 4B**, genes mapped to this GO term in IPF stromal myofibroblasts included numerous genes with established roles in IPF (*1*). Examples of selected strongly prioritized IPF-induced gap GO terms in cell types representing each of the other four compartments are shown in **fig. S10**: cell-cell junction in epithelial AT2 cells (**fig. S10A**); T cell receptor signaling pathway in lymphoid TCD8s (**fig. S10B**); cell leading edge in myeloid monocyte-derived macrophages (**fig. S10C**); and endothelial cell development in CAPAs (**fig. S10D**).

Our therapeutic gap-based strategy was designed to prioritize TFs that were refractory to treatment with approved antifibrotics. Accordingly, we next sought to determine that the dominant position of STAT3 in the untreated IPF-induced pan-lung GRN (**Fig. 3A**) was recapitulated in the IPF-induced gap GRN. To this end, we performed HCT intersection analysis on IPF-induced gap gene sets (i.e., genes induced in untreated IPF but not repressed by either antifibrotic) across the 40 cell types (**table S14**). We next ranked TFs in the IPF-induced gap pan-lung GRN using the same approach as we had previously for the untreated IPF-induced pan-lung network TFs (number of significant footprints and average footprint size). Congruent with its status as a prominent gap TF, STAT3, in addition to TFs with which it has known regulatory relationships, such as NFATC1, TCF12 and FOSL1, had dominant rankings in the pan-lung IPF-induced gap GRN (**Fig. 4C**). We had earlier observed relatively strong effects for approved antifibrotics in the lymphoid compartment (**Fig. 2A** and **table S7**). Consistent with this, TFs with IPF-induced footprint peaks in cells of the lymphoid lineage, such as IRF4, RELA and MTA3 (**table S8**) had appreciably lower rankings in the pan-lung IPF-induced gap GRN (**Fig. 4C**). Of specific relevance to alveolar fibroblasts, STAT3 was by a considerable margin the strongest TF footprint in the IPF-induced gap GRN for this lineage **(Fig. 4D)**.

### STAT3 inhibition reduces the IPF therapeutic gap in precision cut lung slices

Based on our analysis to this point we hypothesized that STAT3 inhibition would reduce the therapeutic gap in IPF. We previously demonstrated the antifibrotic properties in experimental lung fibrosis of the STAT3 inhibitor TTI-101 (*19*), which blocks phosphorylation of STAT3 Y705, a critical requirement for STAT3 activation (*27*). We next used TTI-101 to validate our hypothesis in the PCLS platform, which has recently emerged as a translational model for IPF (*28*). We treated PCLSs with pirfenidone, nintedanib, TTI-101 or vehicle and generated gene expression values for 37 cell types based on distinct marker genes (**fig. S11**). As a preliminary step we again used scCoda to evaluate the effect of TTI-101 on PCLS cell counts (**fig. S12**); full cell counts and summary data are in **table S15**. We observed significantly lower numbers of several cell types in the presence of TTI-101, including CD8 T and natural killer cells, migratory dendritic cells and peribronchial fibroblasts.

We proceeded to generate DEGs (log_2_ FC>0.32 or <-0.32, FDR<0.1) for the 37 PCLS cell types; full SPP-annotated data are in **table S16**. Validating the PCLS analysis, we observed appreciable enrichment of PCLS IPF vs control-induced genes among the corresponding explant untreated IPF-induced genes for 22 of the 37 cell types common to the two systems (**fig. S13A**). Notably, the strongest identity between the two systems was observed for alveolar fibroblasts, the cell type in which we had previously observed the strongest STAT3 explant footprint. Superimposing explant alveolar fibroblast untreated IPF-induced genes on the IPF-induced network in PCLS alveolar fibroblasts (**Fig. 5A**), we observed induction in both systems of many genes with documented roles as both transcriptional drivers and effectors of IPF. Of note, relative to our explant analysis (**Fig. 2A**), the therapeutic gap was larger across the PCLS analysis. This may be attributable to the relatively short duration of exposure to antifibrotics between the two systems, as well as to the fact that some of the PCLS IPF samples were obtained from inviduals who had previously received antifibrotic therapy. Finally, validating the STAT3-inhibitory effect of TTI-101, we observed strong enrichment of STAT3 HCTs among TTI-101-repressed genes across all cell types (**Fig. 5B**), with a peak in alveolar fibroblasts as anticipated.

**Figure 5.**
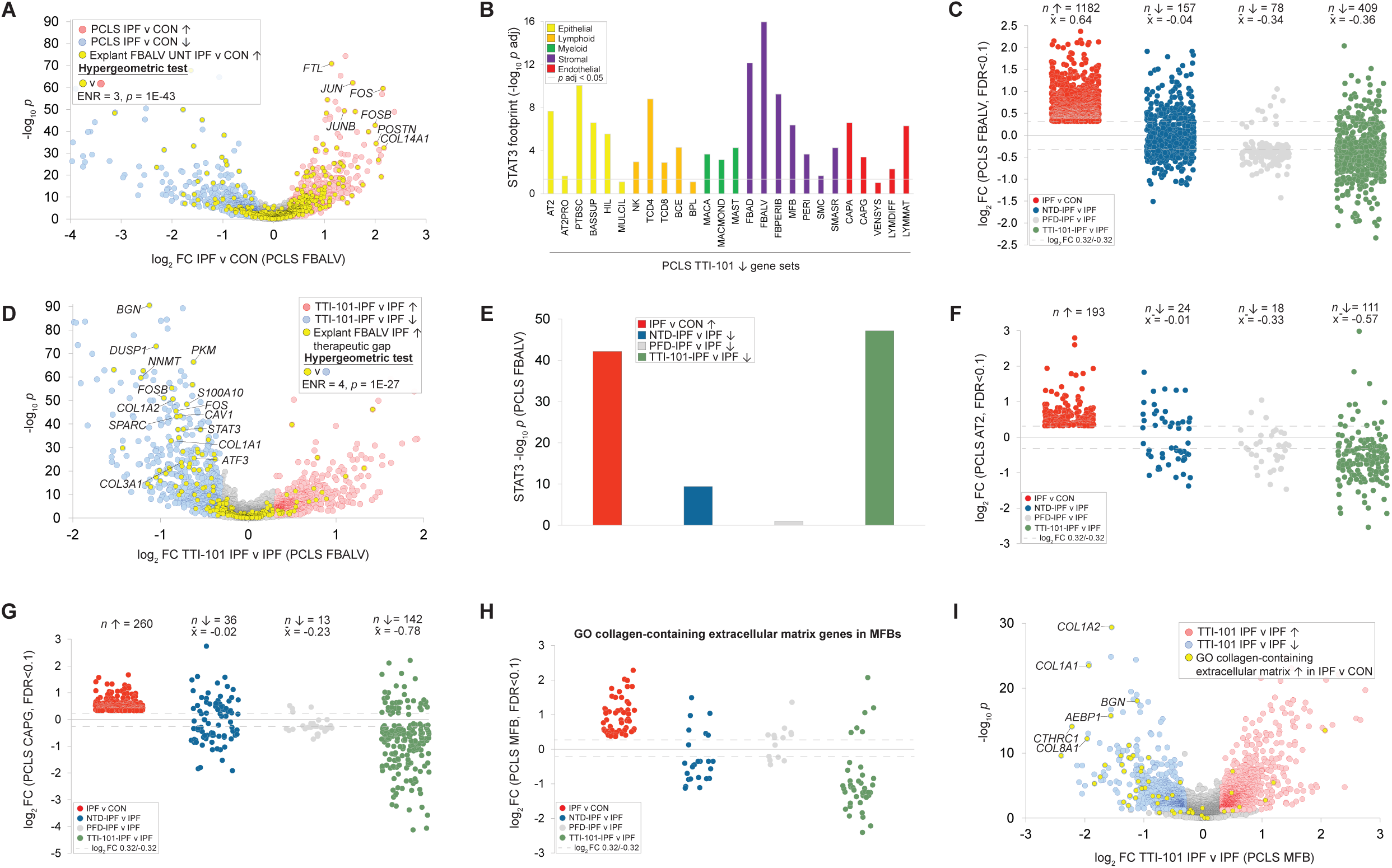
STAT3 inhibition reduces the IPF therapeutic gap in precision cut lung slices. Refer to **table S2** for cell type acronym definitions. **(A)** Volcano plot showing enrichment of explant untreated IPF-induced alveolar fibroblast genes among PCLS IPF-alveolar fibroblast induced genes. **(B)** STAT3 functional footprints of STAT3 among TTI-101-repressed genes across PCLS cell types. **(C)** Manhattan plot comparing the effect on PCLS alveolar fibroblast IPF-induced genes of treatment with nintedanib, pirfenidone and TTI-101. Full data are in **table S16. (D)** Volcano plot showing enrichment of explant alveolar fibroblast untreated IPF-induced therapeutic gap genes among TTI-101-repressed PCLS alveolar fibroblast genes. **(E)** Bar chart comparing STAT3 footprints in PCLS alveolar fibroblast IPF-induced, nintedanib-repressed, pirfenidone-repressed and TTI-101-repressed gene sets. Panels **F-H** are Manhattan plots comparing the effect of treatment with nintedanib, pirfenidone and TTI-101 on **(F)** PCLS AT2 IPF-induced genes. **(G)** PCLS capillary aerocyte IPF-induced genes and **(H)** IPF-induced GO collagen-containing ECM genes in PCLS myofibroblasts. **(I)** Volcano plot showing effect of TTI-101 on specific genes from panel H. Full data are in **table S17**.

We next compared the effect of TTI-101 and approved antifibrotics on IPF-induced genes in PCLS alveolar fibroblasts. We first confirmed that nintedanib (**fig. S13B**) and pirfenidone (**fig. S13C**)-repressed genes from explant alveolar fibroblasts were robustly enriched among respective nintedanib-and pirfenidone-repressed genes in PCLS alveolar fibroblasts, indicating that PCLS recapitulated the *in vivo* effects of approved antifibrotics in this critical IPF cell type. Consistent with our analysis in **Fig. 2B**, we observed strong repression by nintedanib and pirfenidone of ribosomal gene expression in PCLS alveolar fibroblasts (**fig. S13B&C**). Next, comparing the impact of nintedanib, pirfenidone and TTI-101 on the IPF-induced transcriptional program in PCLS alveolar fibroblasts, we found that of 1182 IPF-induced genes, TTI-101 reversed 409, compared with 157 for nintedanib and 78 for pirfenidone (**Fig. 5C**). Moreover, TTI-101 exerted a stronger repressive effect on the IPF-induced program (mean log_2_ FC = - 0.36) than either approved antifibrotic (nintedanib =-0.04, pirfenidone =-0.34). Confirming that TTI-101 reduced the IPF-induced gap in PCLS IPF alveolar fibroblasts, explant alveolar fibroblast IPF-induced gap genes were strongly enriched among TTI-101-repressed genes (ENR = 4, *p* = 1E-27), and included numerous established drivers and effectors of pulmonary fibrosis (**Fig. 5D**). The modest impact of approved antifibrotics on STAT3 function in alveolar fibroblasts, and the extent to which this gap is corrected by treatment with TTI-101, is evident in (**Fig. 5E**), which shows strong STAT3 footprints in IPF-induced and TTI-101-repressed gene sets and relatively weak footprints in nintedanib-and pirfenidone-repressed gene sets.

Although our primary focus was alveolar fibroblasts as the peak of predicted STAT3 activity across the IPF lung, our explant analysis had predicted activation of STAT3 in numerous other prominent cell types (**Fig. 3B**). Accordingly, we were interested to establish whether the strong performance of TTI-101 relative to approved antifibrotics in PCLS alveolar fibroblasts was recapitulated globally across these other cell types. TTI-101 exerted stronger effects than approved antifibrotics in all but three IPF-induced gene sets from PCLS cell types common to all four experimental conditions: untreated, pirfenidone-, nintedanib-and TTI-101-treated (**fig. S13D**). In addition to alveolar fibroblasts, particularly strong relative effects of TTI-101 were observed in two other cell types with predicted peaks of STAT3 activity inferred from the explant analysis, namely AT2 (**Fig. 5F**) and CAPG (**Fig. 5G**) cells. We next compared the effects of nintedanib, pirfenidone and TTI-101 on PCLS IPF-induced genes mapped to the five gap-prioritized GO terms we had previously highlighted (**Fig. 4B** and **fig. S10A-D**). TTI-101 exerted stronger repressive effects than nintedanib or pirfenidone across all five GO terms (**Fig. 5H and fig. S13E-L)**. Reduction of the therapeutic gap by STAT3 inhibition was particularly striking with respect to GO collagen-containing ECM genes in PCLS myofibroblasts: treatment with TTI-101 downregulated 34 IPF-induced members of this term, compared to 18 for nintedanib and three for pirfenidone (**Fig. 5H** and **table S17**). These included members of the collagen chain gene family, in addition to *CTHRC1*, a marker gene of profibrotic fibroblasts (*29*) (**Fig. 5I**).

### STAT3 inhibition disrupts the IPF-induced alveolar fibroblast gene regulatory network

We speculated that the strong antifibrotic effect of TTI-101 in PCLS IPF alveolar fibroblasts relative to approved antifibrotics (**Fig. 5D**) was related to its capacity to reverse STAT3-mediated induction of genes encoding members of the IPF-induced alveolar fibroblast GRN (**Fig. 3G and fig. S8D**). To test this hypothesis in PCLS, we conducted HCT intersection analysis on PCLS alveolar fibroblast IPF-induced genes (**table S18**, column F). Reflecting the strong identity between transcriptional regulatory mechanisms in explant and PCLS alveolar fibroblasts, the top ranked TFs in the explant untreated IPF-induced GRN in this cell type were strongly enriched among the top ranked TFs in its PCLS counterpart (**Fig. 6A**). Similarly, the 23 alveolar fibroblast GRN TF genes that were transcriptionally induced in untreated IPF explants were strongly enriched among PCLS alveolar fibroblast IPF-induced genes (**Fig. 6B**). We next sought to establish that the composition of the PCLS alveolar fibroblast MRN recapitulated that observed in the explant analysis (**Fig. 3J**). Using the same strategy as for the explant MRN (**Fig. 3I**) we performed HCT intersection analysis on the set of 44 PCLS alveolar fibroblast IPF-induced GRN TF genes that were transcriptionally induced in PCLS IPF alveolar fibroblasts per scRNA-Seq (**table S19**). As shown in a scatterplot of the footprints for TFs common to the explant and PCLS IPF alveolar fibroblast MRNs (**Fig. 6C**), we observed a strong correlation (*r* = 0.62, *p* = 9E-08) in footprint size, with STAT3 occupying the top aggregate position across both networks.

**Figure 6.**
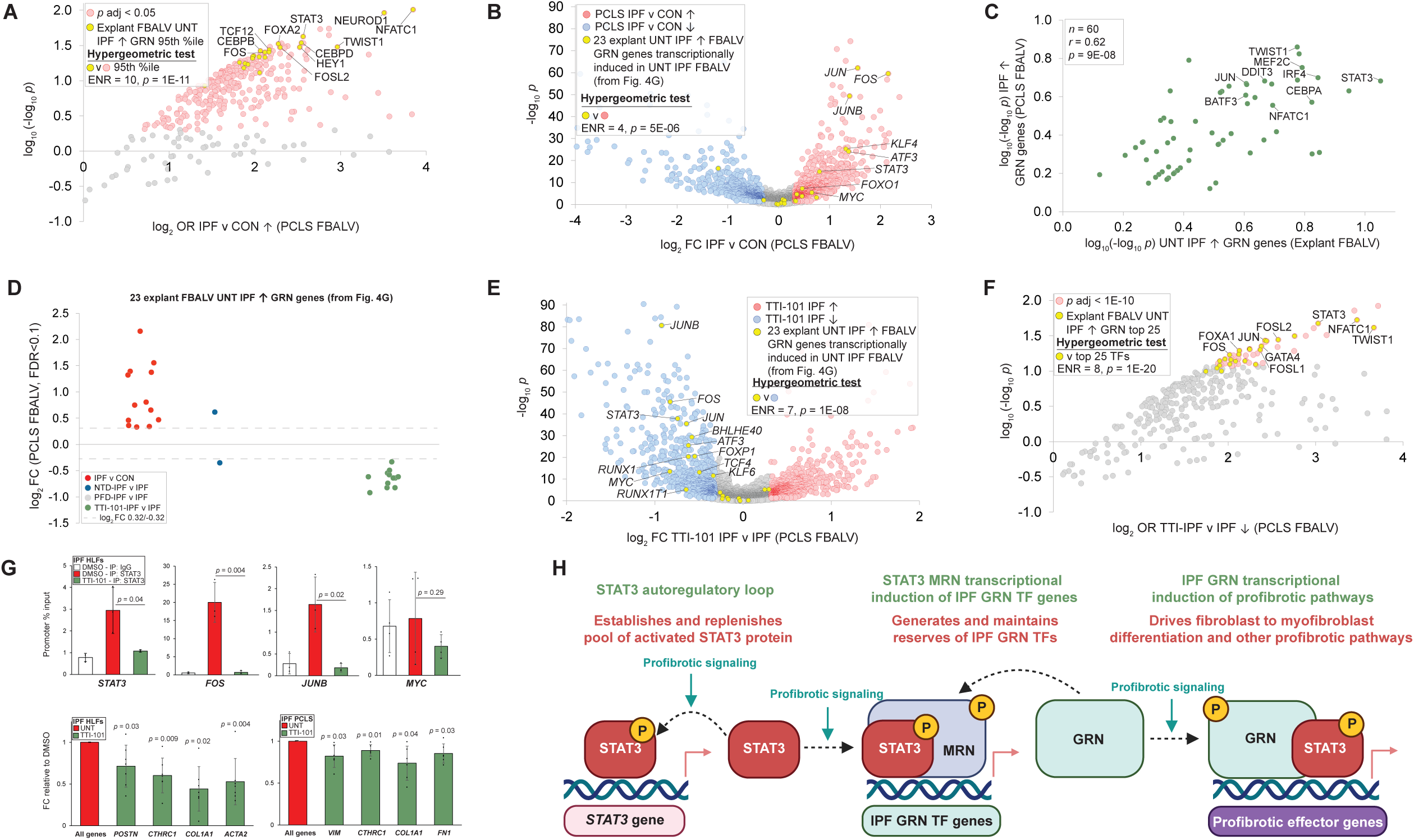
STAT3 inhibition disrupts the IPF alveolar fibroblast gene regulatory network. **(A)** PCLS IPF-induced alveolar fibroblast GRN showing enrichment of top-ranked TFs from the explant alveolar fibroblast untreated IPF-induced GRN. **(B)** Volcano plot of IPF v control PCLS alveolar fibroblast DEGs showing enrichment of genes encoding the 23 untreated IPF-induced explant alveolar fibroblast GRN TFs from **Fig. 3G**. Full data are in **table S16,** column AS**. (C)** Scatterplot comparing-log_10_*p* footprints of explant (*x* axis) and PCLS (*y* axis) alveolar fibroblast IPF MRN TFs. Full MRN data are in **tables S11** (explant) and **table S19** (PCLS). **(D)** Manhattan plot comparing effect of approved antifibrotics and TTI-101 on IPF-induced PCLS FBALV genes encoding the 23 untreated IPF-induced alveolar fibroblast GRN TFs from **Fig. 3G and fig. S9F**. No FDR<0.1 data points existed for the PFD-IPF v IPF contrast. x̄, mean of all data points. **(E)** Volcano plot of TTI-101-IPF vs. IPF PCLS alveolar fibroblast DEGs highlighting TTI-101 repression of the 23 IPF-induced alveolar fibroblast IPF network TF genes. **(F)** TTI-101-repressed PCLS alveolar fibroblast GRN showing enrichment of top TFs from the explant untreated IPF-induced alveolar fibroblast GRN. Full data are in **table S18**. **(G)** Upper panel: STAT3 unloading from promoters of selected FBALV GRN genes was assessed by ChIP-QPCR in IPF HLFs treated with TTI-101 (10 μM) or DMSO. Lower left panel: Q-PCR analysis of genes encoding ECM components in IPF explant-derived HLFs following 24-hour treatment with TTI-101 (10 μM) or DMSO. All target mRNA levels were normalized to *RPLP0*. *p*-values represent comparison to untreated value, i.e. 1. Lower right panel: Q-PCR analysis of genes encoding ECM components in IPF PCLS samples following 72-hour treatment with TTI-101 (10 μM) or no treatment. All target mRNA levels were normalized to *RPLP0*. *p*-values represent comparison to untreated value. i.e. 1. UNT, untreated. See **table S22** for numerical data **(H)** Generalized schematic of STAT3 predicted mechanism of action in IPF alveolar fibroblasts supported by the explant and PCLS analyses. P indicates a phosphorylation event in response to profibrotic signaling. Although GRN and MRN have been depicted as separate entities for clarity, there is extensive overlap in their constituent TFs.

Referring to our explant analysis, we noted that of the 23 alveolar fibroblast untreated IPF-induced GRN TF genes (**Fig. 3G**), only three were repressed by pirfenidone, and seven, most with minor GRN footprints, were repressed by nintedanib (**fig. S14**). Next, we compared the effect of treatment with nintedanib, pirfenidone and TTI-101 on these 23 genes in PCLS. Of the 23 genes from the explant analysis, 13 were IPF-induced in PCLS alveolar fibroblasts (**Fig. 6D**, *p* = 3E-08, hypergeometric test), including *STAT3* and the AP-1 members *FOS*, *JUN* and *JUNB*.

Consistent with their limited effect in the corresponding explant analysis (**fig. S14**), nintedanib treatment reversed only one of the 13 genes, whereas pirfenidone reversed none (**Fig. 6D**). In contrast, treatment with TTI-101 reversed IPF induction of 12 of these 13 genes (*p* = 1E-08, hypergeometric test). **Fig. 6E** visualizes the effect of TTI-101 treatment on the 23 genes in a TTI-101-IPF vs. IPF alveolar fibroblast volcano plot, showing transcriptional repression of genes encoding many IPF GRN TFs with familiar roles in IPF. Reflecting the established autoregulatory capacity of STAT3 (*30*), TTI-101 reversed induction of the *STAT3* gene itself (**Fig. 6E**), whereas treatment with approved antifibrotics had no effect. Given the strong effect of TTI-101 in reversing IPF induction of genes encoding explant alveolar fibroblast GRN TFs in PCLS, we anticipated that TTI-101 would have a profound impact on the functions of proteins encoded by these genes. To evaluate this, we carried out HCT intersection analysis on PCLS alveolar fibroblast TTI-101-repressed gene sets (**table S18, column G**). Reinforcing the robust repressive effect of TTI-101 on alveolar fibroblast IPF GRN function, we observed very strong identity between the top ranked TFs in the explant alveolar fibroblast IPF-induced and TTI-101-repressed GRNs (**Fig. 6F**). As anticipated, STAT3 and AP-1 family members were prominently ranked in the TTI-101-repressed PCLS alveolar fibroblast GRN.

We hypothesized that disruption of the PCLS alveolar fibroblast GRN by TTI-101 was due in part to STAT3 unloading from its own promoter as well as those of genes encoding other components of the GRN. To test this hypothesis in cultured IPF human lung fibroblasts (HLFs), we performed STAT3 chromatin immunoprecipitation targeting the promoter regions of genes encoding *STAT3* and three members of the explant alveolar fibroblast untreated IPF-induced GRN that were among the most transcriptionally responsive to TTI-101 in the PCLS experiment, namely *FOS*, *JUNB* and *MYC*. Supporting our hypothesis, we observed diminished STAT3 loading to all four promoters in TTI-101-treated IPF HLFs (**Fig. 6G**, upper panel). Moreover, Q-PCR analysis of IPF explant-derived HLFs (**Fig. 6G**, lower left panel) and PCLS samples (**Fig. 6G**, lower right panel) confirmed the STAT3 dependence of IPF expression of canonical ECM markers in these cells.

**Fig. 6H** represents a model of STAT3 profibrotic functions in IPF alveolar fibroblasts supported by our explant and PCLS analyses. A STAT3 autoregulatory loop sustained by profibrotic signaling establishes and amplifies a cellular pool of STAT3 that is largely refractory to treatment with approved antifibrotics (left). In response to profibrotic signaling, this pool of STAT3 drives transcriptional induction of genes encoding IPF GRN TFs (middle). STAT3 subsequently co-operates with these TFs in GRNs that recursively drive induction of GRN TF genes (middle) in addition to profibrotic effector pathways, such as fibroblast to myofibroblast differentiation (right) (*26*). In summary, our study positions STAT3 as a prominent transcriptional driver of clinical IPF pathways upon which approved antifibrotics have only a limited impact and, as such, is a priority candidate for the development of novel targeted therapeutics.

### Web resources for data mining of scRNA-Seq and gene regulatory networks

To enable data mining and hypothesis generation from our IPF explant scRNA-Seq and gene regulatory networks, we incorporated them into three open science informatics resources: NDEx, a network visualization resource based on the popular Cytoscape environment (*31*), and our own SPP (*13*) and IPF Cell Atlas (*32*) knowledgebases (**Figure 7**). User documentation on data mining of these resources is in **file S1** and reference 32. Links to these resources for single cell differential gene expression networks and GRNs are in Table 1.

**Figure 7.**
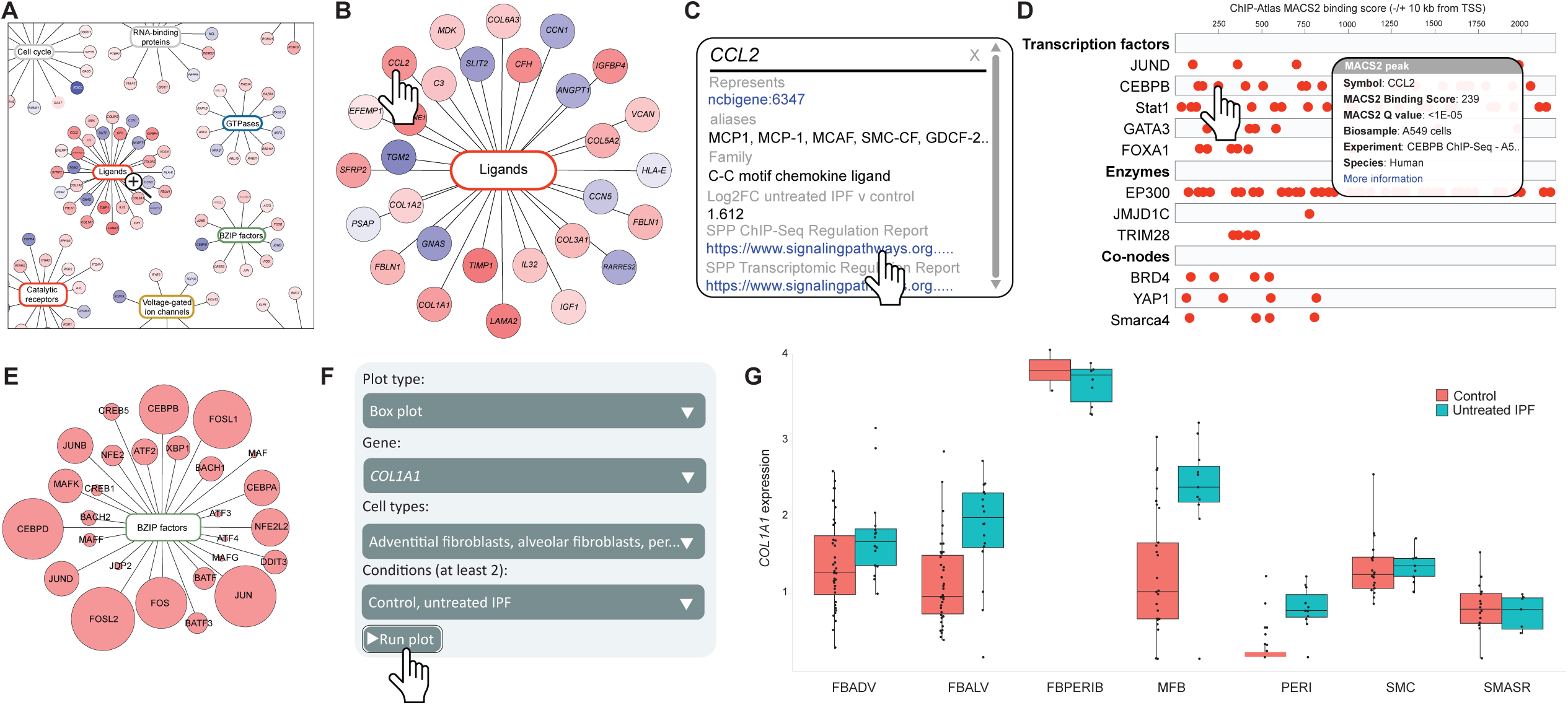
Data mining in the Network Data Exchange, Signaling Pathways Project and IPF Cell Atlas web resources. For user documentation please refer to **file S1** and **ref 32**. **A.** NDEx alveolar fibroblast untreated IPF v control scRNA-Seq network detail showing diversity of gene classes. DEGs in the example Ligands class are shown in **B.** Clicking on a gene (in this example *CCL2*) launches the fold change detail window **(C)**. A link to the NCBI Entrez GeneID record provides comprehensive gene and protein information. Clicking on the SPP link opens the *CCL2* ChIP-Seq Regulation Report in the SPP website **(D)**, which enables the user to identify published experimental evidence for binding of transcription factors, enzymes and co-nodes to the *CCL2* regulatory region. **(E)** Detail of the untreated IPF-induced alveolar fibroblast GRN showing TFs mapped to the BZIP class. Circle size is proportional to footprint-log_10_*p*. **(F)** IPF Cell Atlas Gene Explorer query form. **(G)** IPF Cell Atlas box and whisker plot showing absolute *COL1A1* expression values across stromal cell types in control and untreated IPF samples.

**Table 1.** Links to single cell. RNA-Seq and gene regulatory networks for this study

| Resource | Network type | Link |
| --- | --- | --- |
| NDEx | scRNA-Seq | Untreated IPF v control |
|  |  | Nintedanib IPF v untreated IPF |
|  |  | Pirfenidone IPF v untreated IPF |
|  | Gene regulatory network | Untreated IPF v control-induced |
|  |  | Nintedanib IPF v untreated IPF-repressed |
|  |  | Pirfenidone IPF v untreated IPF-repressed |
| IPF Cell Atlas | scRNA-Seq | IPF Cell Atlas |

## DISCUSSION

Efficient drug discovery demands the prioritization of targets that have broad transcriptional footprints in a given disease state and whose function is not appreciably impacted by existing therapeutics. Although numerous TFs have been implicated in experimental IPF, the disparate nature of the studies from which they emerged has complicated appreciation of their relative transcriptional contribution to the development of clinical IPF. Here we overcame this knowledge deficit by computing a treatment status-segregated single cell RNA-Seq explant analysis against an extensive library of high confidence transcriptional targets for ∼450 human TFs, from which activation of TFs in specific IPF cell types could be inferred and validated. Encouragingly, this pan-lung untreated IPF GRN analysis prioritized numerous TFs that are linked by experimental evidence to the etiology of IPF. Moreover, the prominent ranking of STAT3 in the IPF-induced gap pan-lung GRN indicated that it was for the most part refractory to treatment with approved first-generation antifibrotics. The strong PCLS validation of our prioritization of STAT3 reflects the potential of this approach to discover critical regulatory modules in other cell types of strong relevance to IPF. As such, our study represents the most comprehensive analysis to date towards defining a transcriptional regulatory hierarchy in clinical IPF that can be recapitulated in a translational human model.

The dominant position of STAT3 in the transcriptional hierarchy of IPF alveolar fibroblasts is validated against several independent lines of evidence. Firstly, the ChIP-validated alveolar fibroblast IPF-induced GRN contains numerous proteins, including several members of the immediate early response class, whose encoding genes are known to be regulated by STAT3. Secondly, the alveolar fibroblast MRN bears strong identity with a STAT3 core complex in leukemia, and contains numerous other proteins known to interact with STAT3. Thirdly, the MRN was enriched for TFs encoded by genes shown to have essential, non-redundant roles in both lung fibroblast differentiation and wound healing, including STAT3, SMADs and AP-1 family members. Fourthly, recent experimental studies have predicted a role for STAT3 as a master regulator of fibroblast-to-myofibroblast differentiation and migration (*33, 34*). Moreover, our data align strongly with previous mechanistic studies demonstrating that STAT3 establishes its own regulatory network at the transcriptional and translational levels, through both autoregulation and as a regulator of other TFs (*35*). The prominent position of STAT3 in the IPF alveolar fibroblast MRN may explain why inhibition of a single molecular event, (i.e. phosphorylation of STAT3 Y705) so appreciably inverts the transcriptional landscape of IPF relative to approved antifibrotics in this cell type. Although TFs are historically considered more challenging than conventional pharmaceutical targets, our results indicate that MRN components represent potentially rewarding targets for disease intervention.

Our study has several limitations and caveats. Since the GRN analysis is necessarily restricted to TFs for which ChIP-Seq datasets are archived in public databases, we cannot exclude that TFs not encompassed by our study make significant contributions to IPF pathogenesis. We caution also that for some TFs, GRN rank is impacted by fold change or significance cutoffs for input gene lists, emphasizing the need for robust validation of this analysis. Furthermore, since our study did not include proteomic or metabolomic platforms, it does not address IPF-dysregulated events interrogated by these approaches, nor the extent to which they are impacted by approved antifibrotics. Our PCLS model recapitulated IPF-dysregulated and antifibrotic-reversed transcriptional programs in the majority of IPF cell types. Its failure to do so in all cell types however indicates the need for additional technical refinement of this platform, such as optimizing the duration of drug treatment. It should also be pointed out that our findings on the effect of approved antifibrotics on IPF gene expression in explants are correlative in nature and should be interpreted as such. Finally, it bears mentioning also that STAT3 has been shown to promote alveolar regeneration in AT2 epithelial cells (*36*). As such, untargeted STAT3 inhibition has the potential to adversely impact this and other protective roles of STAT3 in certain cell types.

In conclusion, we applied single cell GRN analysis in a treatment-segregated IPF cohort to identify TFs important in IPF that are weakly targeted by approved antifibrotics. An important outcome of our study was to position STAT3 as a master regulator of the IPF GRN in alveolar fibroblasts and potentially other cell types that drive the pathogenesis of IPF. We anticipate that free access to our dataset through NDEx, SPP and the IPF Cell Atlas knowledgebases will be valuable to investigators seeking to benchmark small molecules of interest against approved antifibrotics, as well as establishing their potential to reduce the therapeutic gap in alveolar fibroblasts and other important IPF cell lineages. More broadly, we envisage that the analytical methodology introduced in our study, involving combined single cell transcriptomics and GRN analysis of a treatment status-segregated cohort, will support the development of novel treatments for other human diseases.

## MATERIALS AND METHODS

### Study design

The aim of our study was to identify novel therapeutic targets in idiopathic pulmonary fibrosis that were not impacted by treatment with approved antifibrotics. The study included explanted lungs from patients with IPF, diagnosed according to current clinical practice guidelines (*37*), which were collected and segregated according to treatment status prior to transplant surgery. This cohort consisted of lungs from 22 untreated, 24 pirfenidone-treated and 28 nintedanib-treated IPF patients. These three cohorts were compared to 64 control lungs from organ donors which lungs were rejected for transplant and procured through an Organ Procurement Organization. The sample size was selected based on available IPF and control tissues cryopreserved in the Chronic Lung Disease Tissue Repository at Baylor College of Medicine (Institutional Review Board H-46823). Before sample collection, written informed consent was provided by all participating subjects. For PCLS experiments, PCLSs were generated from seven IPF lungs and were treated *ex vivo* with either nintedanib, pirfenidone, TTI-101 or their vehicles as described. Single cell RNA sequencing libraries were generated from these lungs. Libraries with low gene counts, high mitochondrial or ribosomal content were excluded from the analysis. Predefined end points included the reversal of IPF-dysregulated gene expression programs using log_2_ FC >0.32 (induction) and <-0.32 (repression) as cutoffs at FDR<0.1.

Females and males of all races, contingent on sample availability, were included in this study. All study participants provided written informed consent approved by the Baylor College of Medicine Institutional Review Board.

### Sample preparation for single cell RNA sequencing

Explanted organs were sliced and washed with cold, sterile phosphate-buffered saline (PBS). Biopsies were cryopreserved in Dulbecco’s Modified Eagle Medium (D-MEM) with 10% DMSO, 10% Fetal Bovine Serum, and 1% Penicillin-Streptomycin-Glutamine, and stored at the Baylor College of Medicine Chronic Lung Disease Tissue Repository (H-46823). After thawing at 37°C, the tissue was minced into small pieces and incubated for one hour at 37°C with D-MEM containing Elastase (30 U/ml, Elastin Products Co. EC134), Recombinant DNAase I (500 U/mg, Roche Diagnostics 04536282001), Liberase (0.3 mg/ml, TM Research Grade 05401127001), and 1% Penicillin-Streptomycin-Glutamine. Digestion was halted with 10% Fetal Bovine Serum. The digested tissue was filtered through a 100-micron mesh and centrifuged at 300g for 10 minutes. Cell pellets were resuspended in red blood cell lysis buffer (VWR International, Radnor, PA) for 3 minutes at 37°C, centrifuged again, and resuspended in MACS medium. For cell concentration and viability assessments, cells were stained with Trypan Blue and counted using a DeNovix Celldrop (DeNovix, USA) or a Countess II Automated Cell Counter (Thermo Fisher Scientific, USA).

### Single cell barcoding, library preparation and sequencing

Single cells were barcoded using the 10X Chromium X single-cell platform, and complementary DNA (cDNA) libraries were prepared following the manufacturer’s protocol (Single Cell 5′ Reagent Kits v2, 10X Genomics, USA). Cell suspensions, reverse transcription master mix, and partitioning oil were loaded onto a single-cell chip with a target of 20,000 cells per library, assuming 100% viability, and then processed on the Chromium X. Reverse transcription occurred within the droplets at 53°C for 45 minutes. cDNA was amplified for 12 cycles using a Bio-Rad C1000 Touch thermocycler. Size selection of cDNA was performed with SPRIselect beads (Beckman Coulter, USA) at a SPRIselect-to-sample volume ratio of 0.6. The resultant cDNA was analyzed using an Agilent Bioanalyzer High Sensitivity DNA chip for quality control. cDNA was fragmented with a proprietary enzyme blend, followed by end-repair and A-tailing at 65°C for 30 minutes. Double-sided size selection was conducted using SPRIselect beads.

Sequencing adaptors were ligated to the cDNA at 20°C for 15 minutes. cDNA was then amplified with a sample-specific index oligo as a primer, followed by another round of double-sided size selection using SPRIselect beads. Final libraries were analyzed on an Agilent Bioanalyzer High Sensitivity DNA chip for quality control. Libraries were sequenced on the Illumina NovaSeq 6000 system. Cell Ranger (10X Genomics, USA) was used to generate FASTQ files from sequencing data. Adapter sequences were trimmed from the FASTQ data, and reads were demultiplexed, aligned, and counted. 10x quality control metrics are provided in **table S20**.

### Cell type annotation and differential gene expression

Single cell lung data was mapped using 10x Genomics Cell Ranger v7.0.1 onto the human genome reference provided by 10x Genomics. Doublet detection was performed using scrubblet (*38*). Data were processed using the Python Scanpy library (*39*). We performed guided cell annotation using CellTypist (*40*) and references provided by HLCA (https://data.humancellatlas.org/) in addition to our previous aberrant basaloid cell type reference (*5*). UMAP plots were generated using Scanpy. Objects were converted to the Seurat format using the zellkonverter package in R. Cell type marker plots were generated using Seurat (*41*). Differential gene expression was generated using MAST (*42*) with significance achieved at FDR-adjusted p-value<0.1 and fold change exceeding ±log_2_ 0.32 (1.25x induced or repressed). Relative differences in abundance of cell types within specific compartments were derived using ChiSquare as implemented in the R statistical system.

### Chromatin immunoprecipitation

Chromatin immunoprecipitation (ChIP) was performed using the Pierce Magnetic ChIP Kit (Cat #26157) (Thermo Scientific) according to the manufacturer’s instructions. Briefly, approximately 4×106 cells were crosslinked with 1% formaldehyde for 10 minutes at room temperature, followed by quenching with 1X glycine. Cells were lysed to isolate nuclei, and chromatin was digested using Micrococcal Nuclease (MNase) at 37C for 15 minutes to obtain fragments ranging from 200 to 1000 bp. Chromatin was incubated with normal Rabbit IgG (Cell Signaling Technology cat #3900S) or anti-STAT3 (CST cat #30835S) antibody overnight at 4°C. Complexes were captured using Protein A/G Magnetic Beads, washed, and eluted. Crosslinks were reversed at 65°C with NaCl and Proteinase K, followed by DNA purification via spin columns. Promoter specific primers are shown in **table S21**.

### Quantitative PCR

Total RNA was isolated with RNeasy kit (QIAGEN) and cDNA was synthesized from RNA (0.15-1 μg) using iScript Reverse transcription supermix for reverse transcription (RT) PCR (Bio-Rad Laboratories, Hercules, CA, USA). RT-PCR, with SYBR Green Master Mix (Bio-Rad Laboratories, Hercules, CA, USA), was performed using the CFX96 Touch Real-Time PCR Detection System (Bio-Rad Laboratories, Hercules, CA, USA). The relative quantity of target mRNA was calculated by use of the CT method, or 2–ΔΔCT, as described, and normalized by use of RPLP0 as an endogenous control. Target gene primers are shown in **table S21**.

### Primary human lung fibroblasts

Primary human lung fibroblasts (HLFs) were derived from IPF patients who were subjected to lung transplantation. Fibroblasts were cultured in complete media (DMEM; Corning) containing 10% FBS (Corning), 100 IU of penicillin and 100 μg/ml streptomycin (Corning), 292 μg/ml L-glutamine (Corning) in humidified incubators at 37°C and 10% CO2. Once fibroblasts reached desired confluency of 80%, complete media was replaced with low serum media containing 1% FBS (Corning), 100 IU of penicillin and 100 μg/ml streptomycin (Corning), 292 μg/ml L-glutamine (Corning) in humidified incubators at 37°C and 10% CO2 and treated with TTI-101 to a concentration of 10 μM or equal volume DMSO for 24 hours.

### Statistical analysis

For single cell RNA-Seq, differential gene expression was generated using MAST (*42*) with significance achieved at FDR-adjusted p-value<0.1 and fold change exceeding ±log_2_ 0.32 (1.25x induced or repressed). DEGs determined at FDR<10% aligned well with previously published fibrotic transcriptional metrics. Relative differences in abundance of cell types within specific compartments were derived using ChiSquare as implemented in the R statistical system. For Gene Ontology and HCT intersection analysis, a hypergeometric test was formulated with the following parameters: *N* = population size, *k* = number of successes in the sample, *s* = sample size and M = number of successes in the population. Significance was set at *p* < 0.05. For ChIP and Q-PCR, data are presented as mean ± SEM with numerical data provided in the supplementary material. For comparisons between two groups, we used ratio paired t-test or paired/unpaired t-test as suitable. Statistical significance was defined as *p*<0.05.

## LIST OF SUPPLEMENTARY MATERIALS

### Supplementary Materials and Methods

#### Supplementary figures

Figures S1-S14

#### Supplementary tables

Tables S1-S22

#### Other supplementary files

File S1. Instructions for navigating NDEx and SPP web resources.

---

**Fig. S14.**
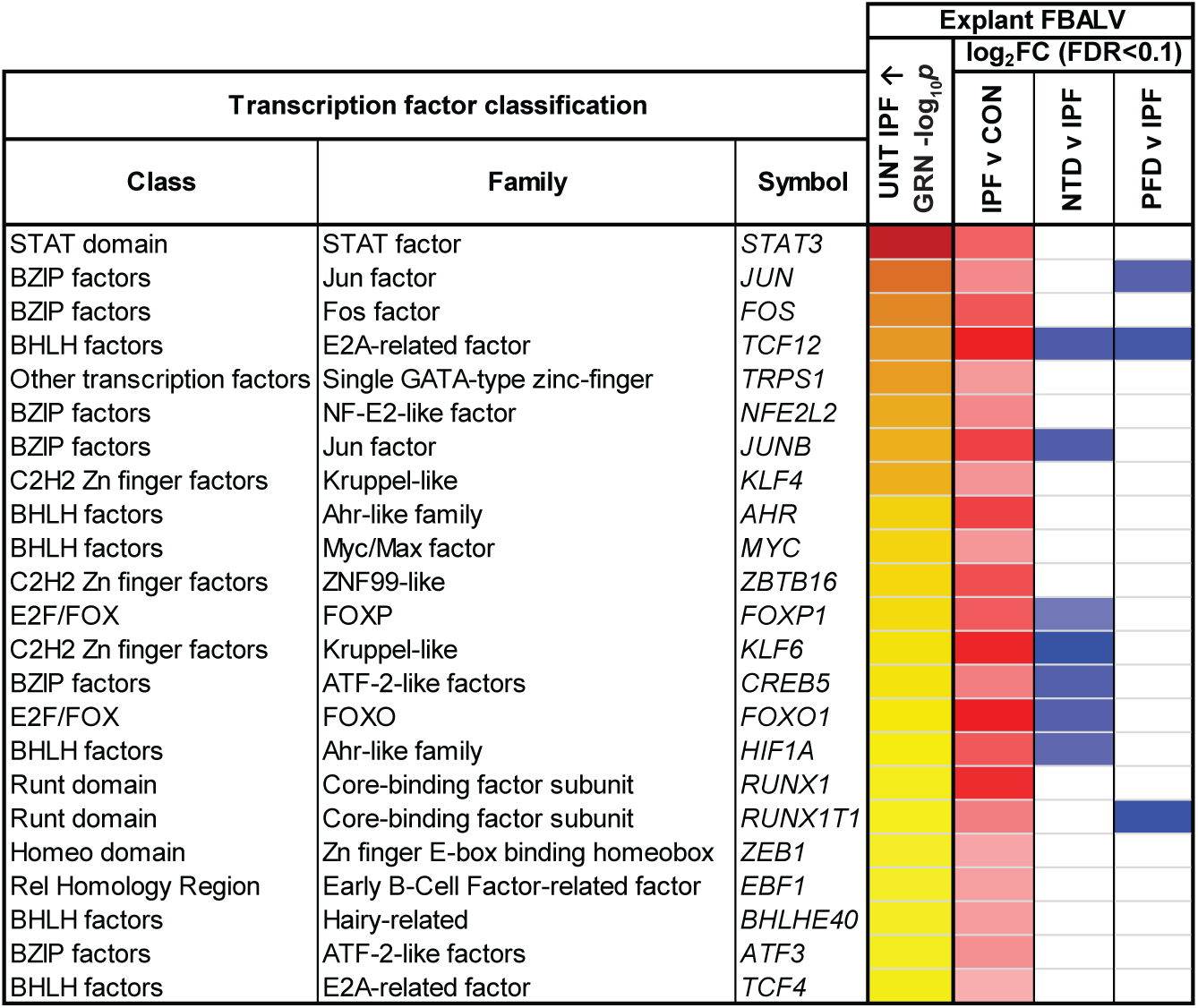
Effect of nintedanib and pirfenidone on the 23 untreated IPF-induced explant alveolar fibroblast genes encoding IPF GRN TFs. Full data are in tables S5 (scRNA-Seq) and table S8 (GRN).

## Precision cut lung slice (PCLS) model

Explanted organs were collected under the Chronic Lung Disease Tissue Repository at Baylor College of Medicine (H-46823) with a combined warm and cold ischemia time of less than 12 hours. A complete lobe was dissected from each explant and inflated through the airway using a 60 ml syringe and a 10 Fr catheter with a solution of DMEM, 2% pre-heated low melting point agarose (UltraPure LMP Agarose, Invitrogen, USA), and 1% PSG. The agarose-filled lobe was cooled on wet ice (approximately 4°C) to allow solidification. The lobe was then sliced longitudinally and cored using a 10 mm diameter punch biopsy (P1025, Acu Punch, USA). Each core was subsequently glued to a coring tool, and the remaining space was filled with the previously described agarose solution. Cores were sliced into 500 µm PCLS using a VF-300-0Z Microtome (Precisionary Instruments, USA). Freshly sliced PCLS were immediately placed on a dish filled with DMEM/F12 + 1% FBS chilled on ice. Each PCLS was placed in a well of a 24-well plate and incubated overnight (37°C, 5% CO₂) in 1.5 ml of medium composed of DMEM/F-12 + GlutaMAX 1X (Gibco, Catalog # A41920-01), 1% Penicillin-Streptomycin (Corning, Catalog #30-002-Cl), and 1% FBS (Corning, Catalog # 35-011-CV). After overnight incubation, the medium was changed to 1.5 ml of treatment medium composed of DMEM/F-12 + GlutaMAX 1X (Gibco, Catalog # A41920-01), 1% Penicillin-Streptomycin (Corning, Catalog #30-002-Cl), and 0.1% FBS (Corning, Catalog # 35-011-CV). After 48 hours of incubation, the medium was changed, and six unique PCLS were treated with the previously described treatment medium and one of the following treatment conditions: 1 µM nintedanib (Sigma-Aldrich, SML2848-25MG), 500 µM pirfenidone (R&D Systems, 1093), or 10 µM TTI-101 (C188-9; Tvardi Therapeutics, USA), their respective vehicles, or without additional treatment. The PCLS were incubated for 72 hours with a medium change at 36 hours. After incubation, PCLS were collected and digested for downstream single-cell RNA sequencing analysis as described above.

## Gene Ontology analysis and hierarchical clustering

Gene Ontology (GO) enrichment analysis for all three GO subontologies was carried out for each of the 40 cell types from the untreated IPF vs control contrast using a universe derived from the Bioconductor org.Hs.eg.db (v. 3.15.0) human annotation package and the enrichGO function from the clusterProfiler (v. 4.4.4) BioC analysis package in R (v. 4.2.1). The 40 cell types from the untreated IPF vs control contrast were hierarchically clustered using GO terms which were significantly enriched (FDR < 0.05) in at least 6 or more of the 40 cell types.

Heatmaps were generated using the Bioconductor pheatmap (v. 1.0.12) analysis package with default settings for Euclidean row and column clustering distance and complete clustering method.

## Analysis of Lung Tissue Research Consortium expression array

To process expression array data for GSE32537, we utilized the log_2_ summarized and normalized array feature expression intensities provided by the investigator and deposited in GEO. These data are available in the corresponding “Series Matrix File(s)”. The full set of summarized and normalized sample expression values for samples labeled with a final diagnosis of “control” or “IPF/UIP” were extracted and processed in the statistical program R (v. 4.2.1). The IPF/UIP samples were further subcategorized into three disease severity groups based on the reported forced vital capacity pre-bronchodilator % predicted values (Mild = FVC > 72%, Medium = FVC < 72% and > 50%, Severe = FVC < 50%). To calculate differential gene expression for experimental contrasts, we used the linear modeling functions from the Bioconductor limma (v. 3.52.3) analysis package. Initially, a linear model was fitted to a group-means parameterization design matrix defining each experimental variable (control, mild, medium, and severe). Subsequently, we fitted a contrast matrix that recapitulated the sample contrasts of interest, in this case IPF/UIP severe vs Control, IPF/UIP medium vs Control, and IPF/UIP mild vs Control, producing fold-change and Benjamini-Hochberg FDR corrected significance values for each array feature present on the Affymetrix Human Gene 1.0 ST Array. The current Bioconductor array annotation library was used for annotation of array identifiers.

## High confidence transcriptional target intersection analysis

We first retrieved processed gene lists from ChIP-Atlas (*43*), in which genes are ranked based on their mean MACS2 peak strength across available archived ChIP-Seq datasets in which a given TF is the IP antigen. We then mapped the IP antigen to its category, class and family, and organized the ranked lists into percentiles to generate human TF ChIP-Seq consensomes.

Genes in the 95th percentile of a given TF consensome were designated HCTs for that TF and used as the input for the HCT intersection analysis using the Bioconductor GeneOverlap analysis package implemented in R (v. 1.32.0). *p* values were adjusted for multiple testing using the method of Benjamini & Hochberg to control the false discovery rate as implemented with the p.adjust function in R, to generate *p* adjusted values. The universe used in the HCTI intersection analysis was set at an estimate of the total number of transcribed (protein and non-protein-coding) genes in the human genome (43,000). Evidence for a transcriptional regulatory relationship between a TF and the gene set of interest was represented by a larger intersection between the gene set and HCTs for a given node than would be expected by chance after FDR correction (*p* adj < 0.05). For scRNA-Seq regulatory networks, cutoffs for gene lists were log_2_FC>0.32 (induced) or <-0.32 (repressed) at FDR < 0.1. For microarray data, cutoffs for gene lists were log_2_FC>0.32 (induced) or <-0.32 (repressed) at *p* < 0.05. These “discovery-mode” cut-offs were based upon a desire to maximize the number of input genes for the GRN analysis, which in turn allowed us to confidently call a larger number of members of the corresponding GRN.

## Immunohistochemistry

Immunohistochemical staining was performed on formalin-fixed, paraffin-embedded lung explant tissue sections from three IPF cases and three non-fibrotic control subjects matched for age, sex, race, and smoking status. Staining for periostin (POSTN; Abcam, Cat# ab215199), fibronectin (FN1; Abcam, Cat# ab2413), and collagen I (COL1A1; Novus Biologicals, Cat# PA2140-2) was performed by HistoWiz (Brooklyn, NY, USA). Whole-slide digital images were generated by HistoWiz, and representative regions from control and IPF lungs, including preserved parenchyma and honeycomb areas, were selected by a board-certified pathologist for visualization. Semiquantitative analysis was performed in QuPath v0.7 using positive pixel classification, and the percentage of DAB-positive tissue area was calculated using marker-specific thresholds.

