## Supplementary Figures for "Single cell transcriptomics in a treatment-segregated cohort exposes a STAT3-regulated therapeutic gap in idiopathic pulmonary fibrosis"

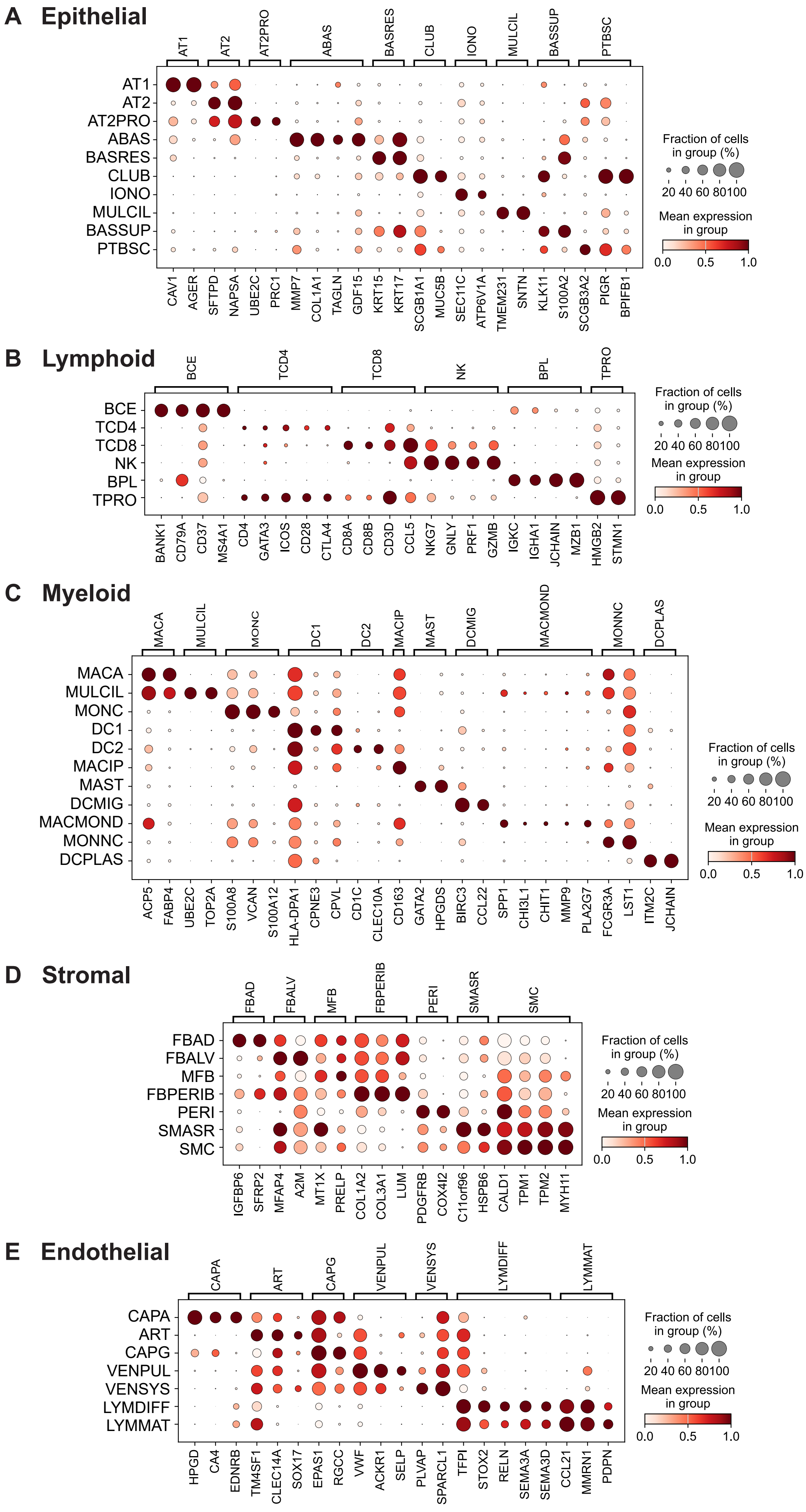


**Fig. S1. Marker genes for lung explant cell types.**

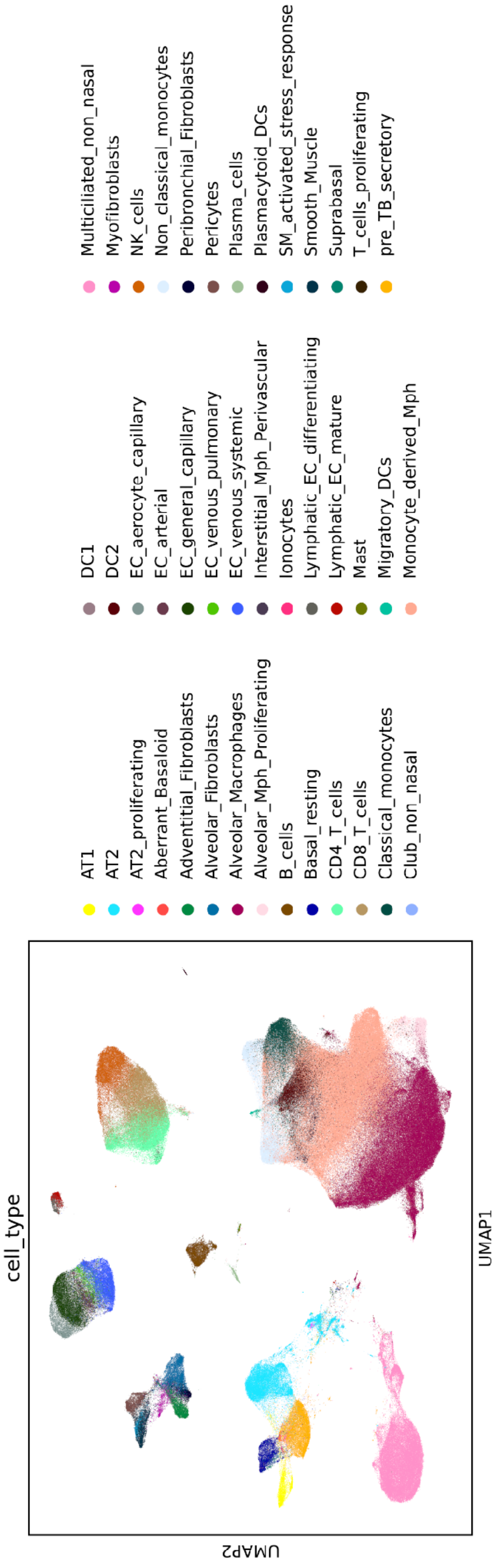


**SCVI Explant**

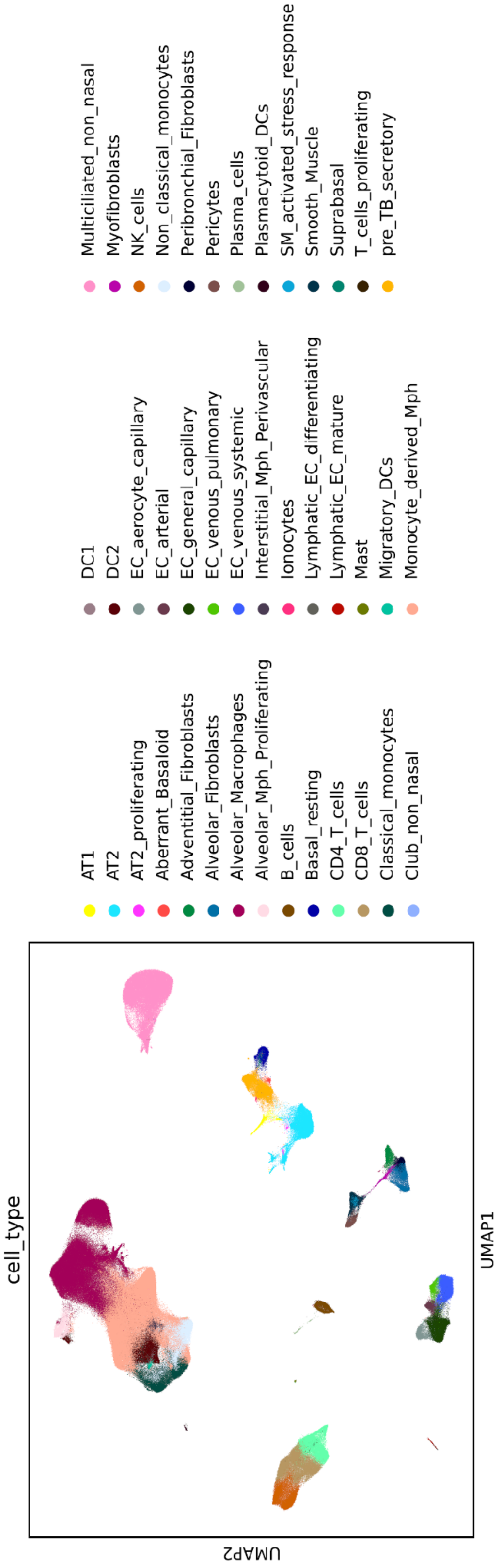


**BBKNN Explant**

**Fig. S2.** Comparison of SCVI and BBKNN Uniform Manifold Approximation and Projection (UMAP) representation of explant cell types.

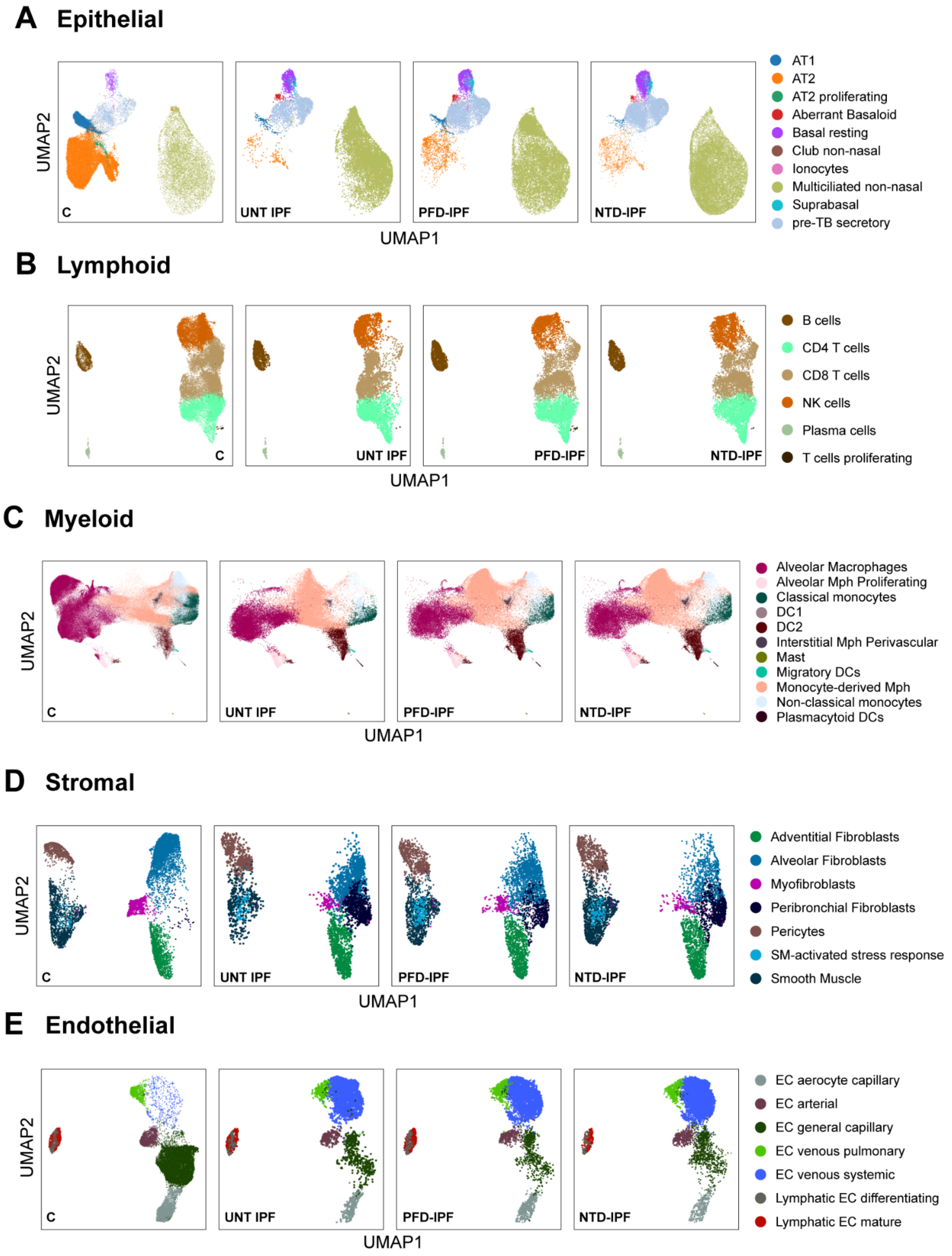


**Fig. S3.** Uniform Manifold Approximation and Projection (UMAP) representation of cells in the epithelial **(A)**, lymphoid **(B)**, myeloid **(C)**, stromal **(D)** and endothelial **(E)** compartments from each of the explant treatment groups. C, control; UNT IPF, untreated IPF; PFD-IPF, pirfenidone-treated IPF; NTD-IPF, nintedanib-treated IPF.

### EXPLANTS: UNTREATED IPF v CONTROL

**For all plots: x axis = untreated IPF v control $\log_2$FC FDR<0.1 unadjusted for covariates; y axis = untreated IPF v control $\log_2$FC (no FDR cutoff) adjusted for covariates**

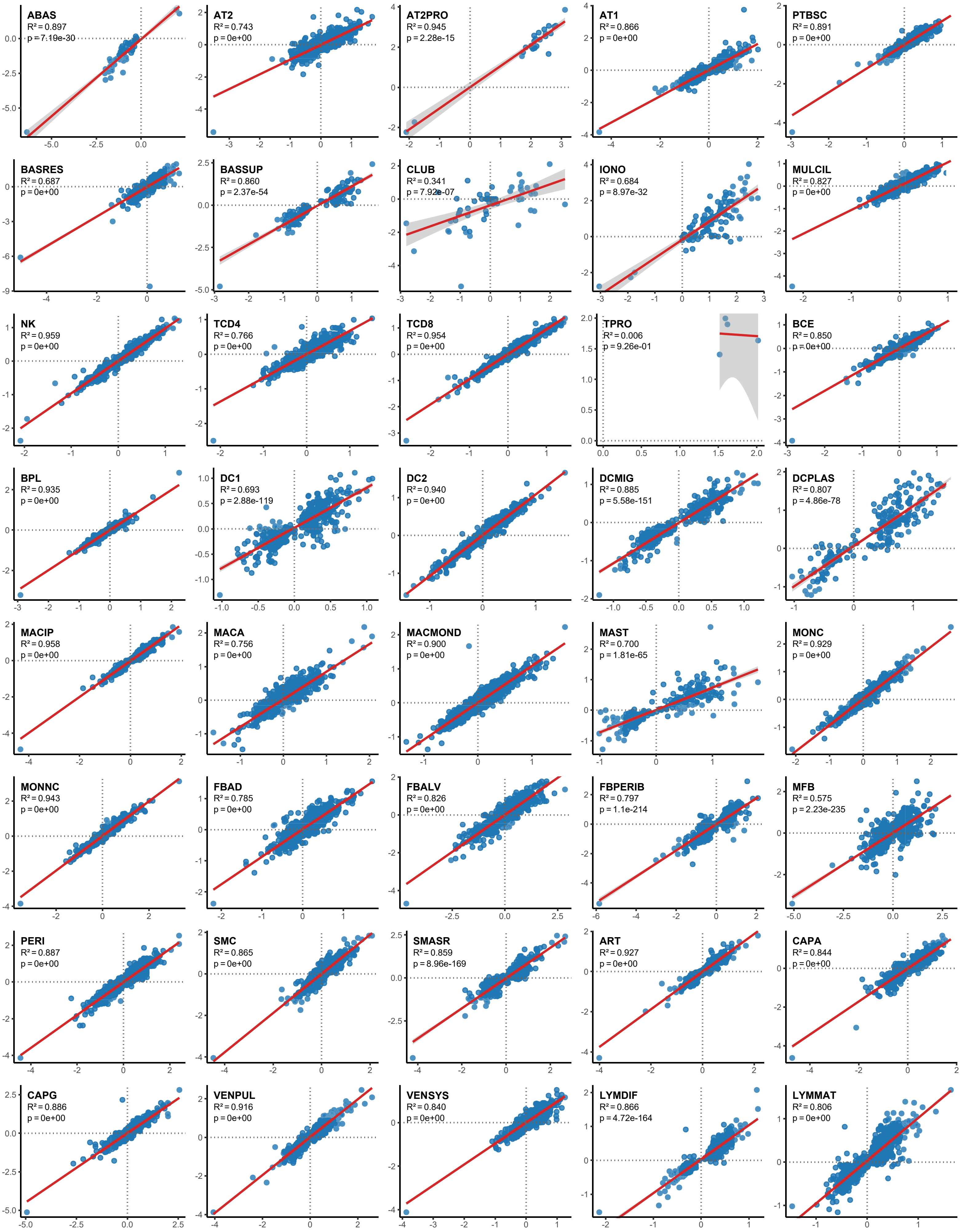

### EXPLANTS: NINTEDANIB-IPF v UNTREATED IPF

**For all plots: x axis = nintedanib IPF v untreated IPF $\log_2$FC FDR<0.1 unadjusted for covariates; y axis = nintedanib IPF v untreated IPF $\log_2$FC (no FDR cutoff) adjusted for covariates**

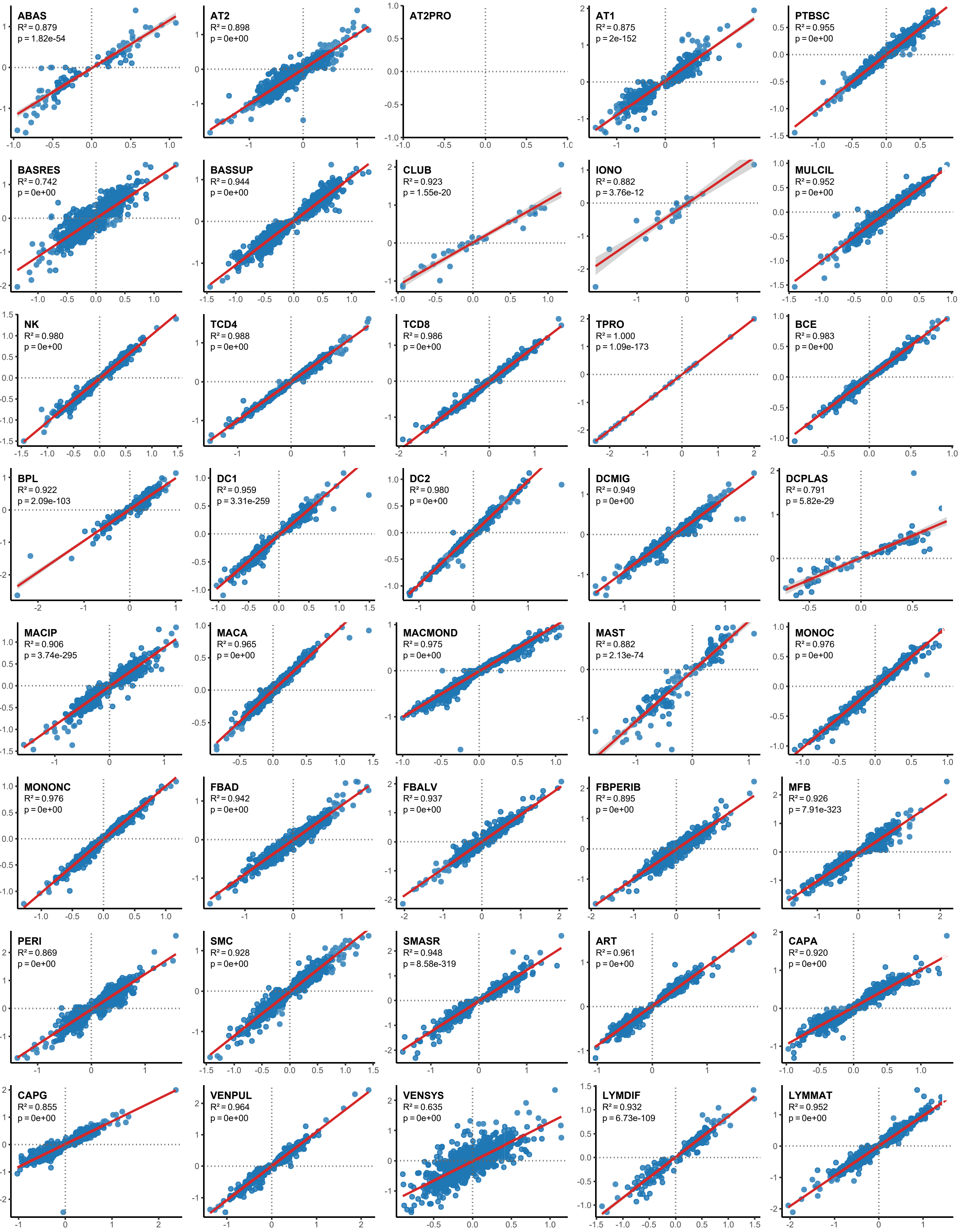

### EXPLANTS: PIRFENIDONE-IPF v UNTREATED IPF

**For all plots: x axis = pirfenidone IPF v untreated IPF $log_2FC$ FDR<0.1 unadjusted for covariates; y axis = pirfenidone IPF v untreated IPF $log_2FC$ (no FDR cutoff) adjusted for covariates**

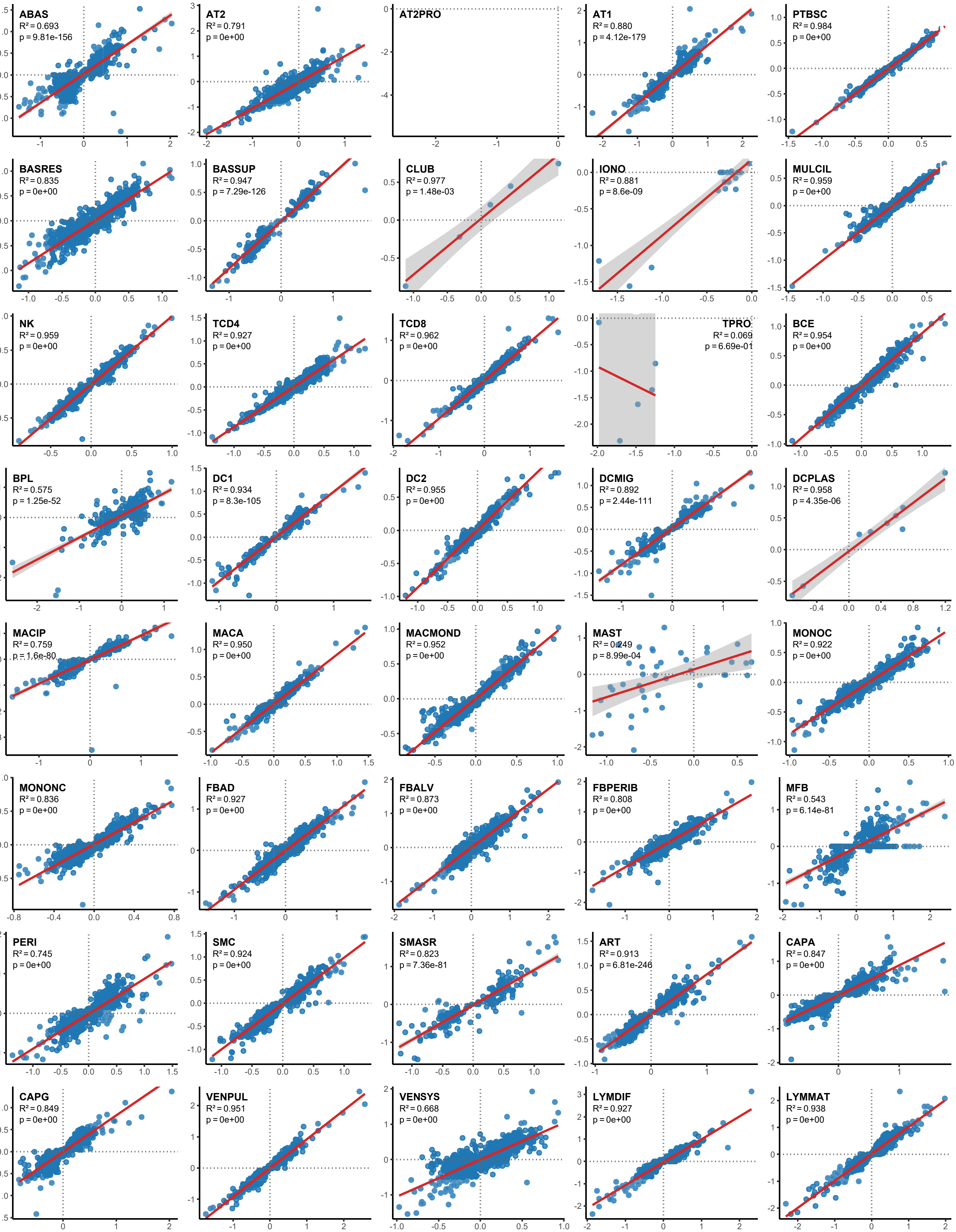

**Fig. S4. Evaluation of confounding effect of age, sex and smoking status covariates in explants.** L2FC values for untreated IPF v control, nintedanib IPF v untreated IPF and pirfenidone IPF v untreated IPF explants were adjusted as described. For each cell type Pearson's correlation analysis was applied to L2FC values from the unadjusted dataset that had an FDR <0.1 against the gene's adjusted L2FC value without FDR cutoff. With the exception of a couple of select cell type/contrast combinations (e.g. club cells in untreated v control) we observed strong alignment between adjusted and unadjusted L2FC values across all cell type/contrast combinations, indicating the absence of significant confounding by age, sex and smoking status covariates.

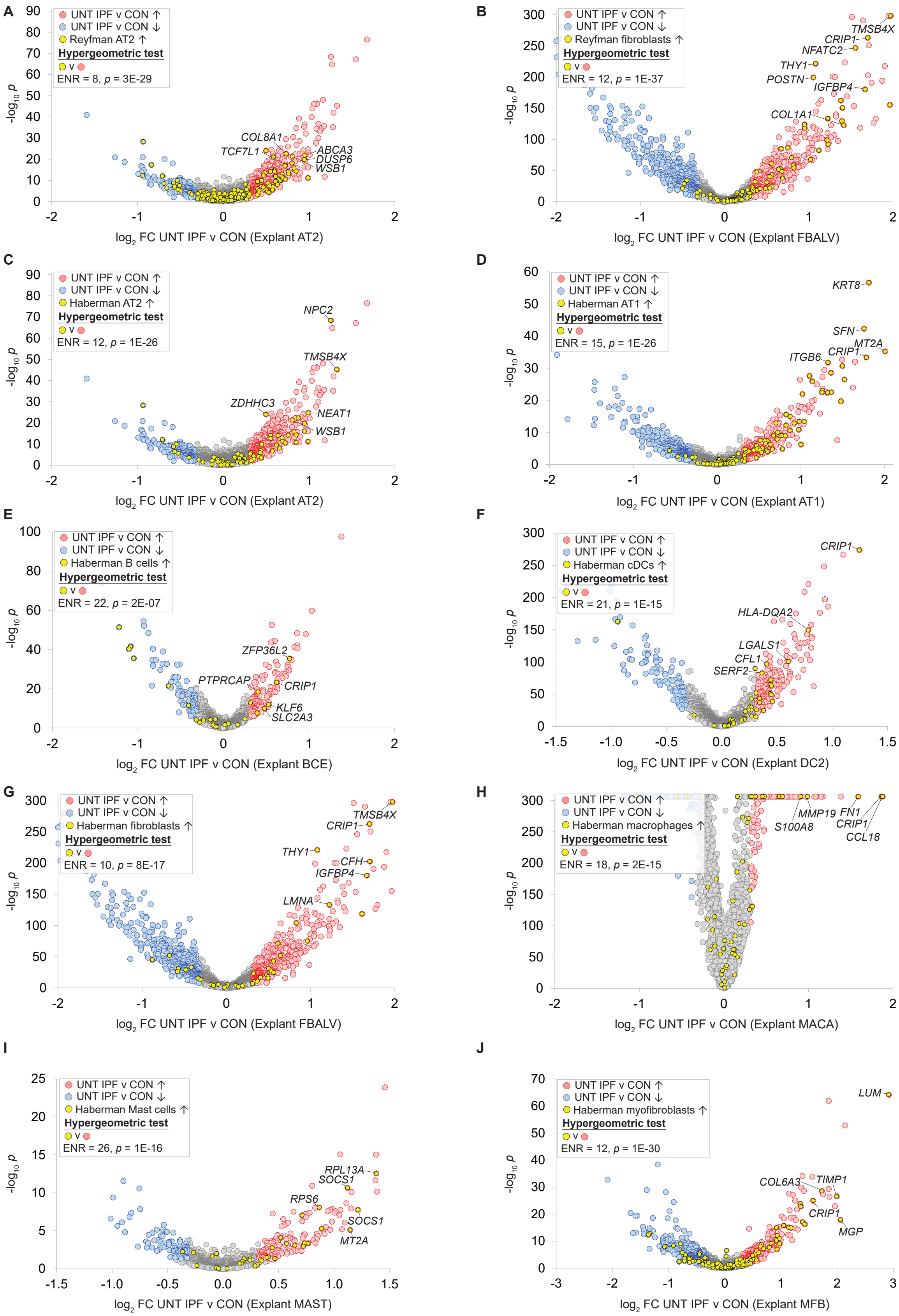


**Fig. S5. Alignment of untreated IPF vs control explant differential gene expression from the current study with published Reyfman and Haberman IPF single cell atlases.** Panels A-J represent untreated IPF vs control volcano plots for the indicated explant clusters from the current study. Overlaid on each are IPF-induced genes from comparable clusters retrieved from the supplementary data for the indicated study: Haberman et al. (6) and Reyfman et al (8). Enrichment statistics indicate broad alignment between the datasets with respect to IPF-induced gene sets. **(A)** our AT2 vs Reyfman AT2; **(B)** our FBALV vs Reyfman fibroblasts; **(C)** our AT2 vs Haberman AT2; **(D)** our AT1 vs Haberman AT1; **(E)** our BCE vs Haberman B cells; **(F)** our DC2 vs Haberman cDCs; **(G)** our FBALV vs Haberman fibroblasts; **(H)** our MACA vs Haberman macrophages; **(I)** our MAST cells vs Haberman mast cells; **(J)** our MFB vs Haberman myofibroblasts. Note that since the current study adopted Human Lung Cell Atlas annotations that the Reyfman and Haberman studies predated, comparison between identical clusters was not possible. Moreover, unlike our study, the Reyfman and Haberman studies combined treated and non-treated IPF subjects in the same IPF cohort.

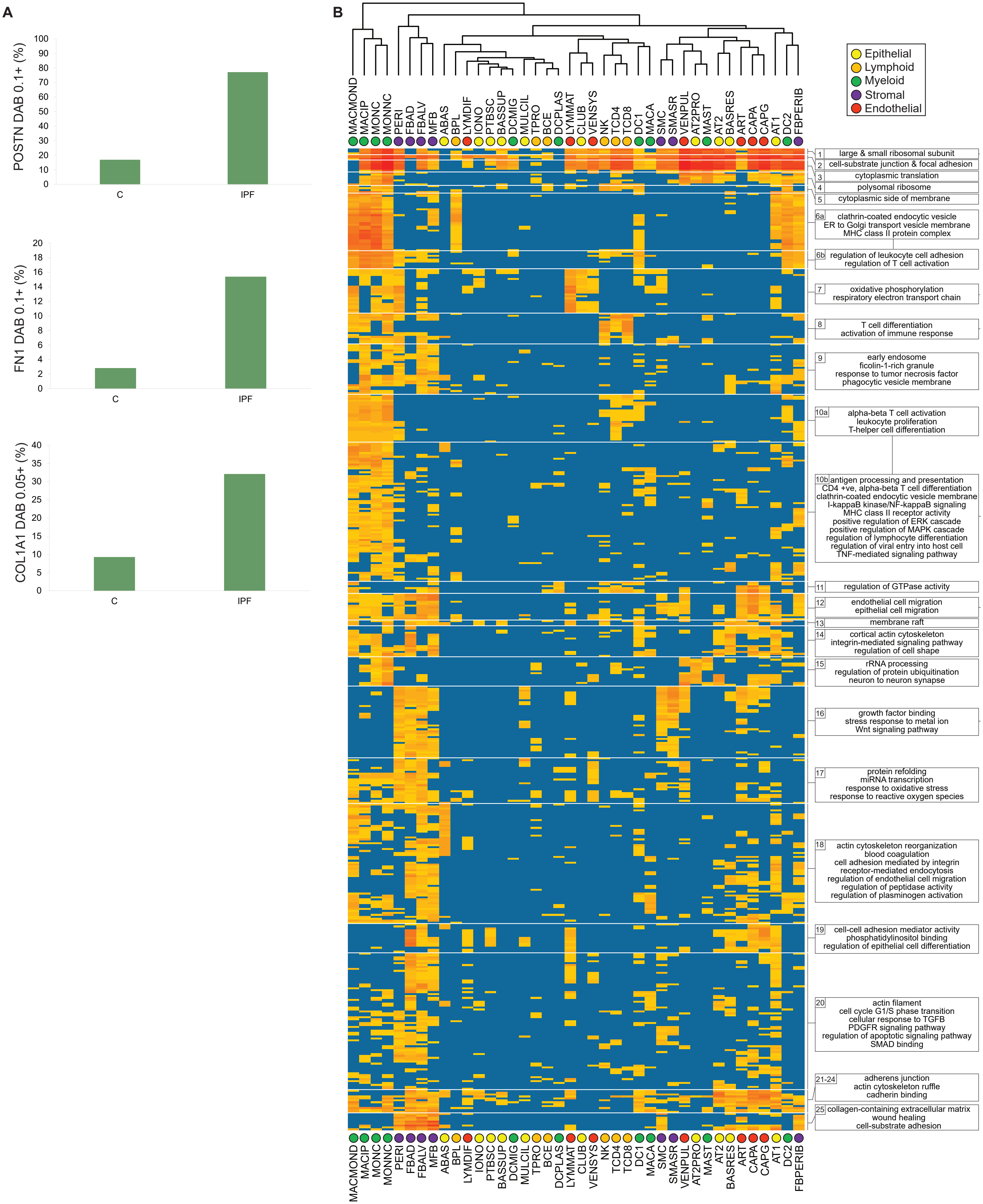


**Fig. S6.** **(A)** Semiquantitative analysis of immunohistochemical validation of untreated IPF v control induction of *POSTN*, *FN1* and *COL1A1* from scRNA-Seq analysis. C, control; IPF, IPF preserved parenchyma **(B)** Full untreated IPF-induced GO heatmap for top 25 clades. See text for details and table S6 for full dataset.

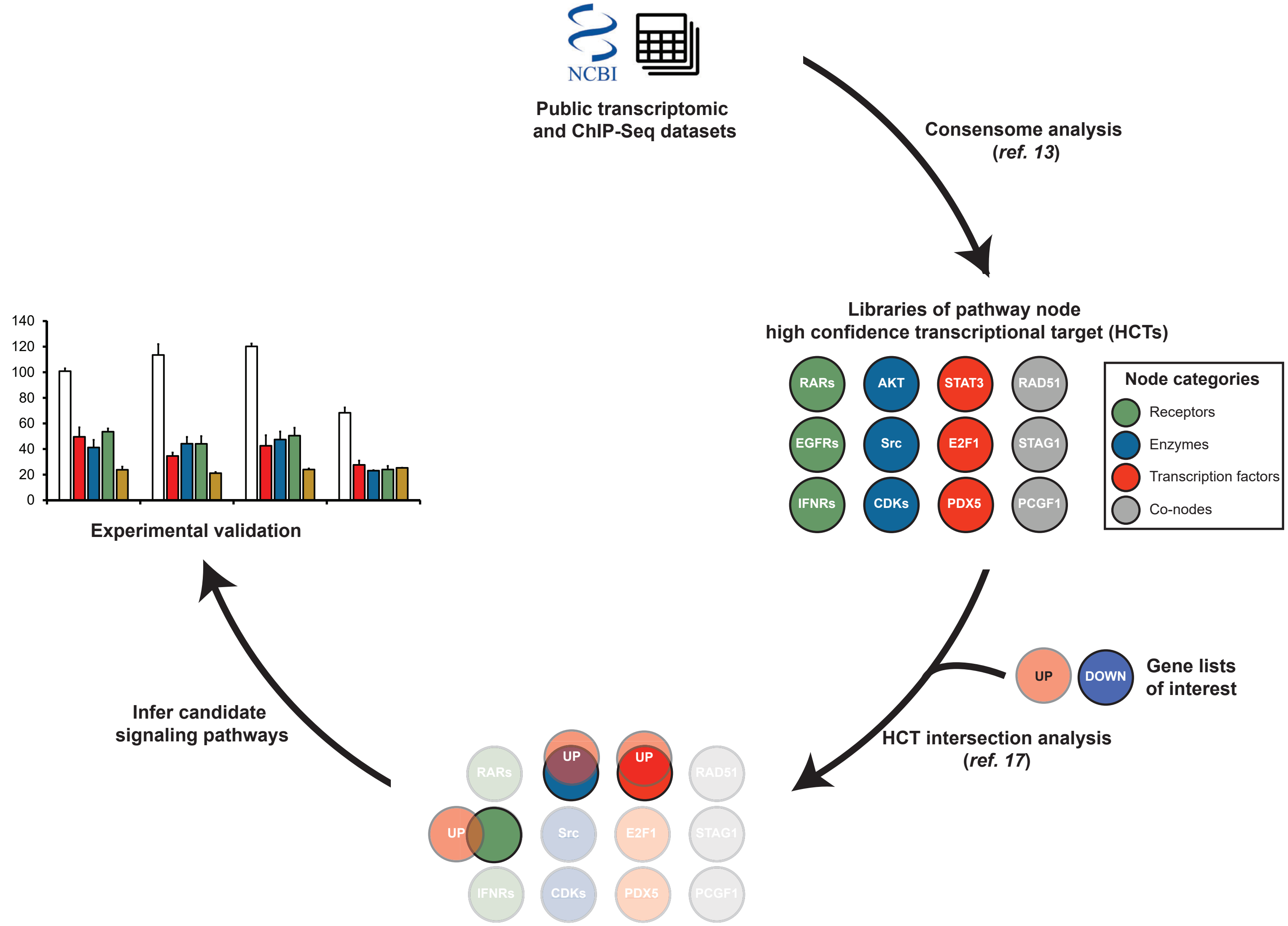


**Fig. S7. Overview of the high confidence transcriptional target (HCT) intersection analysis platform for generating gene regulatory networks**. **(A)** Consensome analysis (*13*). Over 10,000 public transcriptomic or ChIP-Seq experiments were mapped to their pathway node or biosample of study. To enable prediction of pathway node-gene target transcriptional regulatory relationships, we generated consensus 'omics signatures, or consensomes, which ranked genes based on measures of their significant differential expression or promoter occupancy in experiments mapped to a specific node family. **(B)** HCT intersection analysis (*17*). The 95th percentile of each consensome was defined as high confidence transcriptional targets (HCT) for a specific node or node family. To predict gene regulatory networks for a given gene set, a hypergeometric test was used to compute the overlap between HCTs and the gene sets of interest. **(C)** Validation. Loss- and gain-of-function experiments are designed to validate the predictions generated by the GRN analysis.

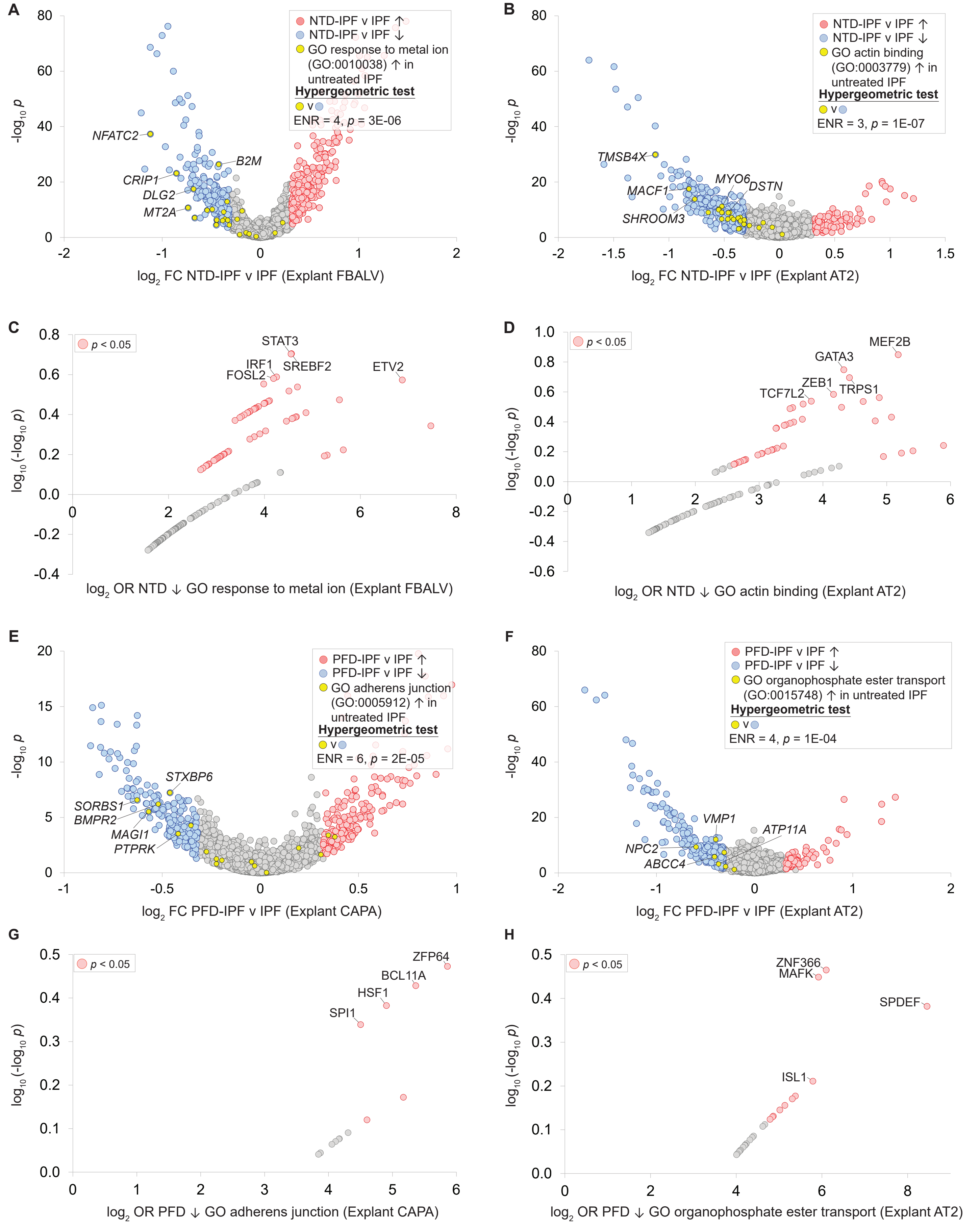


**Fig. S8. Effects of approved antifibrotics on specific GO term-mapped genes in selected cell types.** Volcano plots in panels **A** & **B** show enrichment of untreated IPF-induced genes mapping to selected GO terms in nintedanib-repressed genes in the indicated cell types. Gene regulatory network plots in panels **C & D** indicate TFs predicted to be repressed by nintedanib in panels **A & B**, respectively. Volcano plots in panels **E** & **F** show enrichment of untreated IPF-induced genes mapping to selected GO terms in pirfenidone-repressed genes in the indicated cell types. Gene regulatory network plots in panels **G & H** indicate TFs predicted to be repressed by pirfenidone in panels **E & F**, respectively.

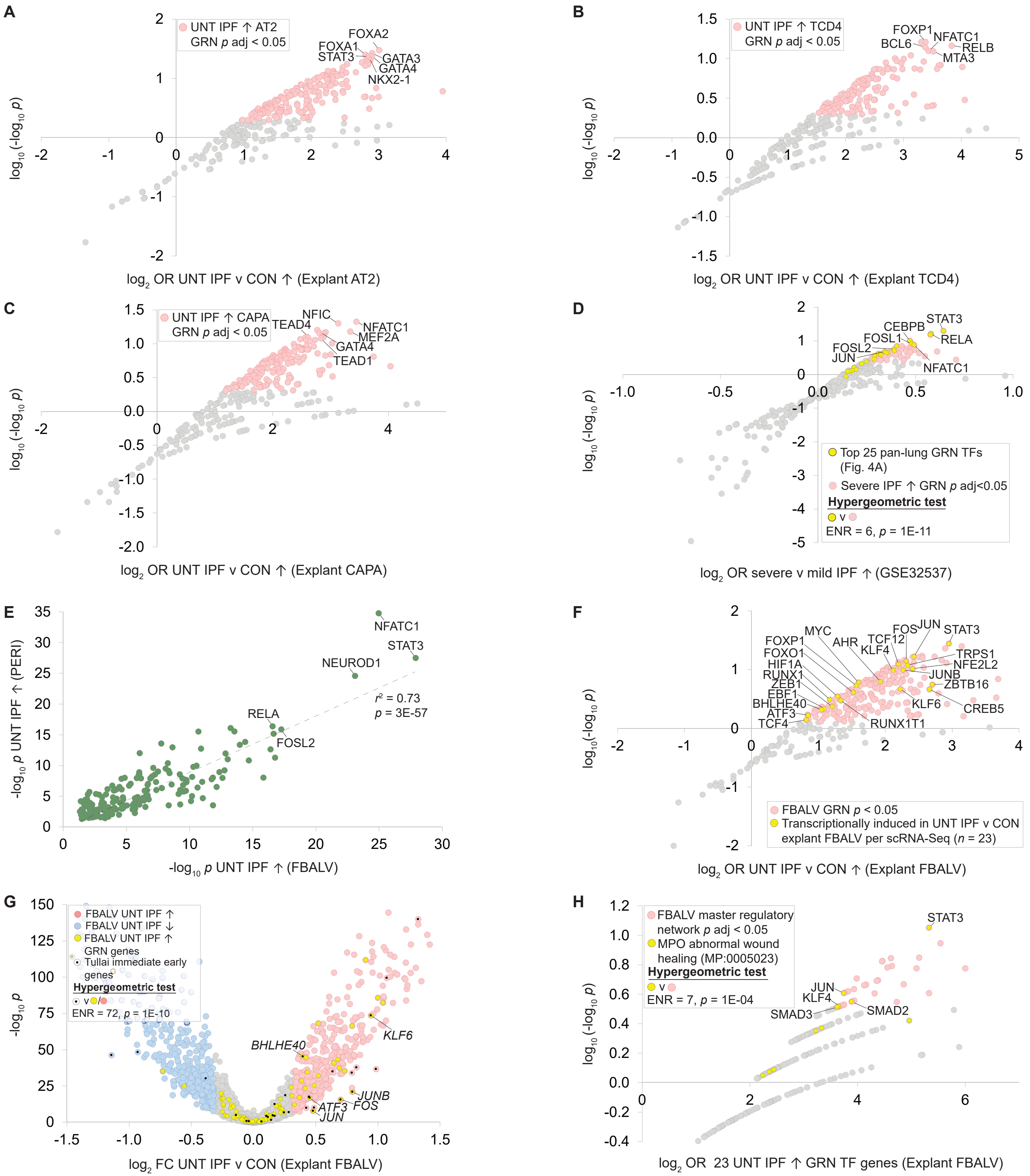


**Fig. S9. Refer to table S2 for cell type abbreviations.** Scatterplots **A-C** represent untreated IPF-induced gene regulatory networks for AT2 cells **(A)**, TCD4s **(B)** and CAPAs **(C)**. TFs with the strongest and most significant footprints within the gene set of interest are distributed towards the top right of the plot. **(D)** GRN plot showing enrichment of the top 25 untreated IPF-induced pan-lung GRN TFs among the top ranked TFs in the severe v mild IPF GRN derived from Lung Tissue Research Consortium (LTRC) dataset GSE32537. Full data are in **table S11**. **(E)** Scatterplot comparing TF -log10*p* footprints among untreated IPF-induced genes in alveolar fibroblasts (x axis) and pericytes (y axis). **(F)** Untreated IPF-induced explant alveolar fibroblast GRN showing TFs that are transcriptionally induced in untreated IPF. **(G)** Enrichment in the set of 23 untreated IPF scRNA-Seq-induced alveolar fibroblast GRN TF genes of a set of experimentally-defined immediate early genes (IEGs). IEGs were retrieved from table S5 in Tullai et al. (*22*). **(H)** Enrichment in the alveolar fibroblast MRN of human orthologs of TFs encoded by mouse genes mapping to the Mammalian Phenotype Ontology (MPO) term "abnormal wound healing (MP:0005023). Genes mapping to this MPO term were retrieved from the Mouse Genome Informatics MouseMine resource (www.mousemine.org).

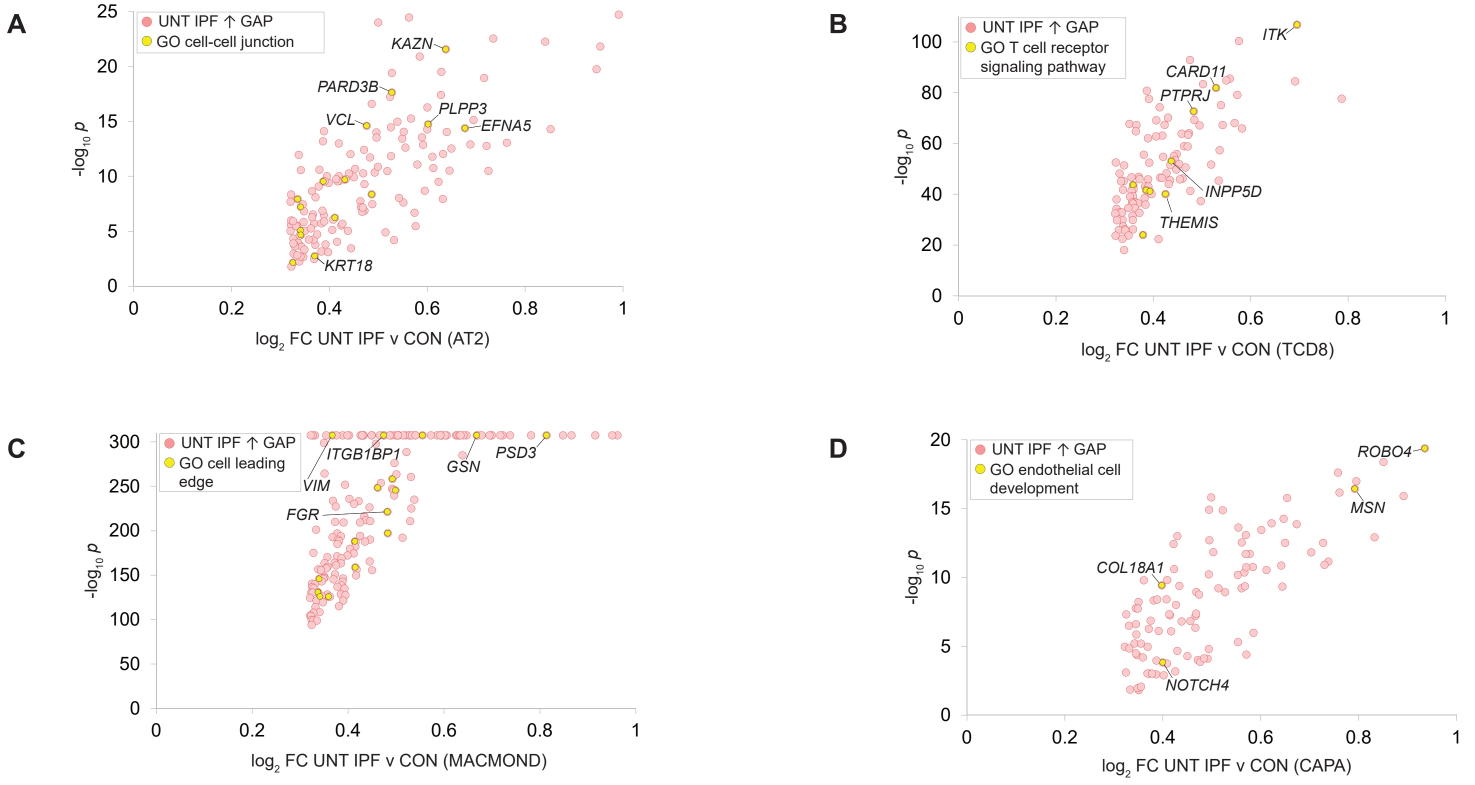


**Fig. S10.** Induction in untreated IPF vs control of genes mapped to the following gap-prioritized GO terms: **(A)** cell-cell junction in AT2 cells; **(B)** T cell receptor signaling pathway in TCD8s; **(C)** cell leading edge in MACMONDs; **(D)** endothelial cell development in CAPAs.

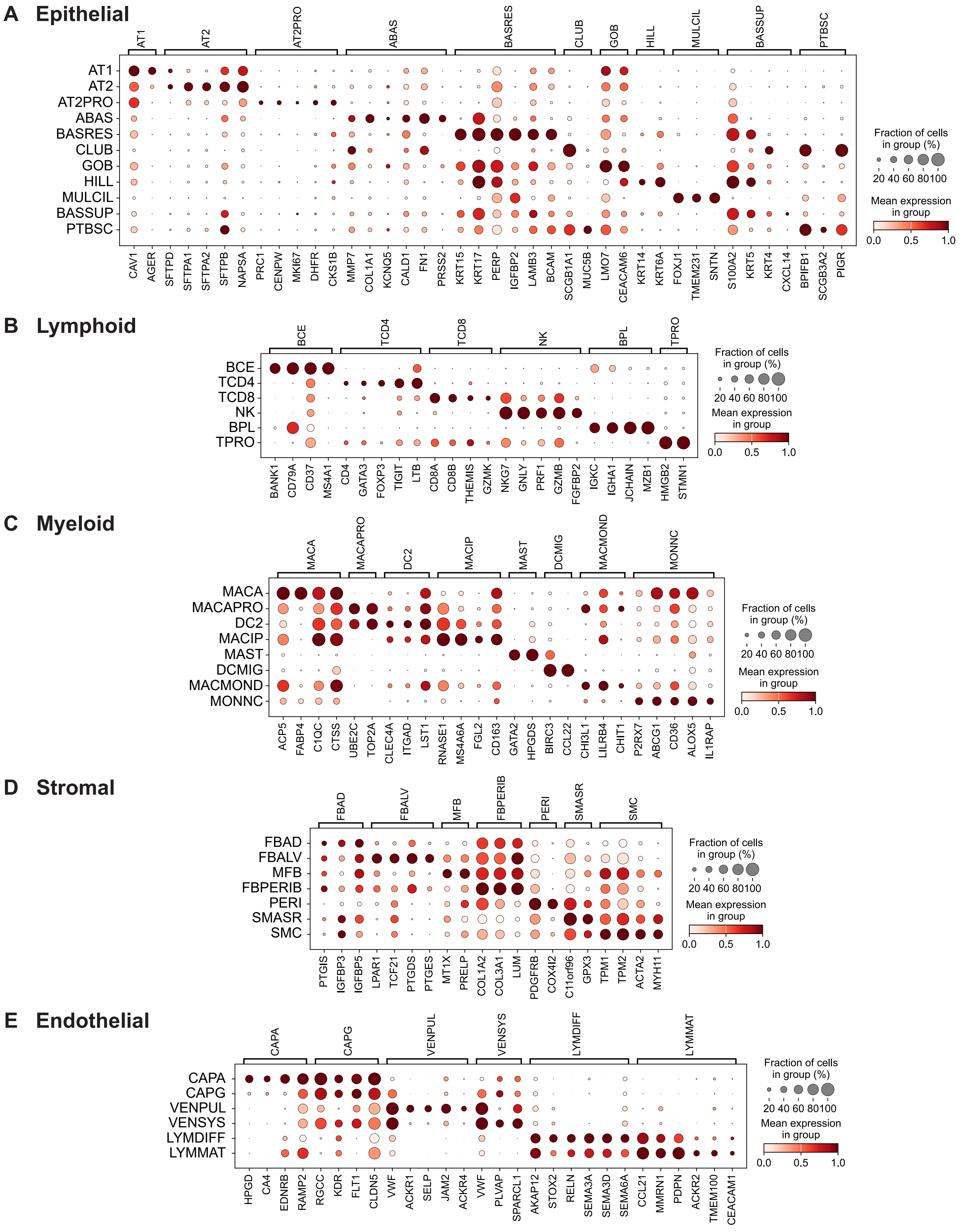


**Fig. S11.** Marker genes for PCLS cell types in epithelial **(A)**, lymphoid **(B)**, myeloid **(C)**, stromal **(D)** and endothelial **(E)** compartments.

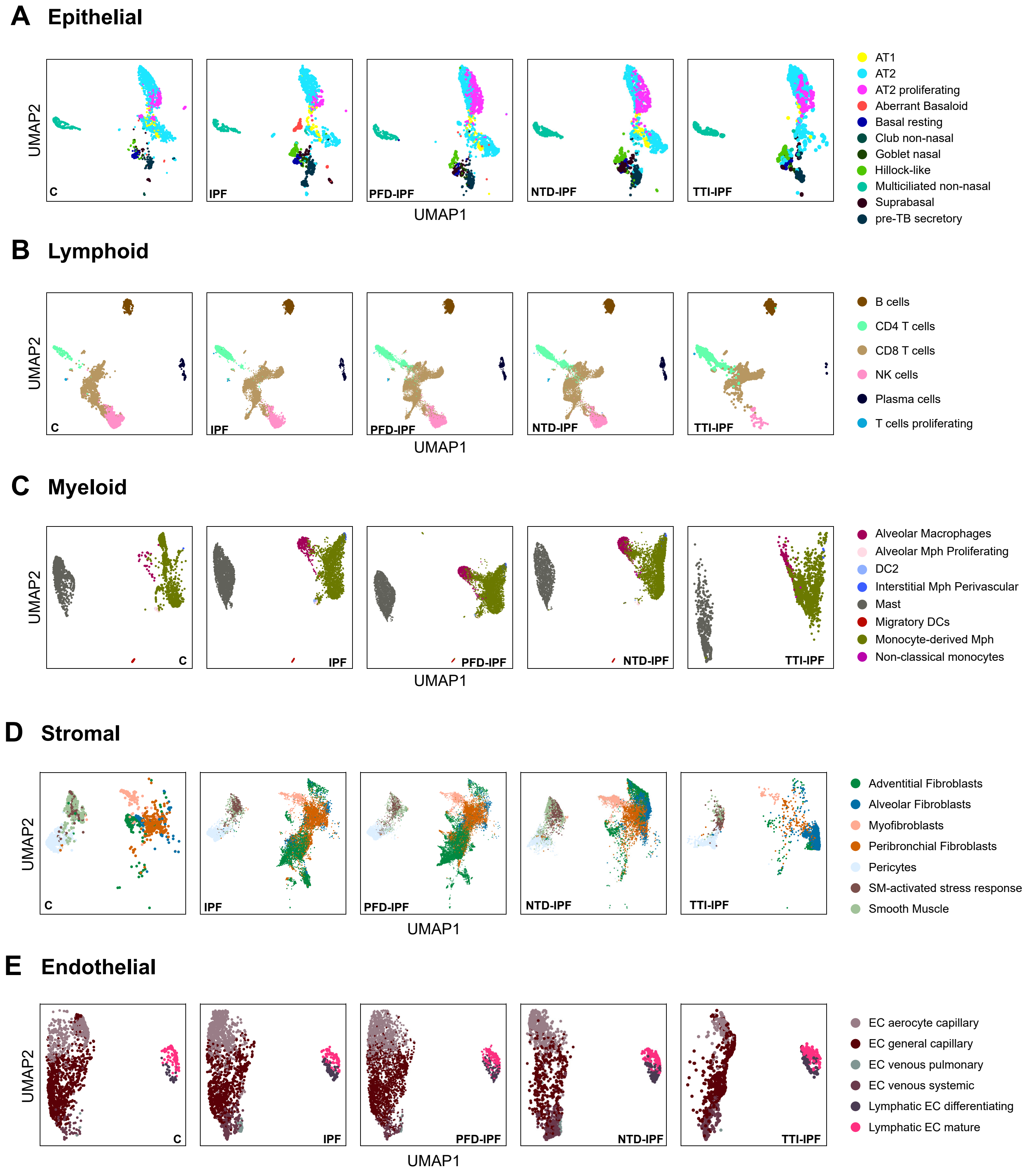


**Fig. S12.** UMAP representation of cells in the epithelial **(A)**, lymphoid **(B)**, myeloid **(C)**, stromal **(D)** and endothelial **(E)** compartments from each of the PCLS treatment groups. C, control; IPF, IPF; PFD-IPF, pirfenidone-treated IPF; NTD-IPF, nintedanib-treated IPF; TTI-IPF, TTI-101-treated IPF.

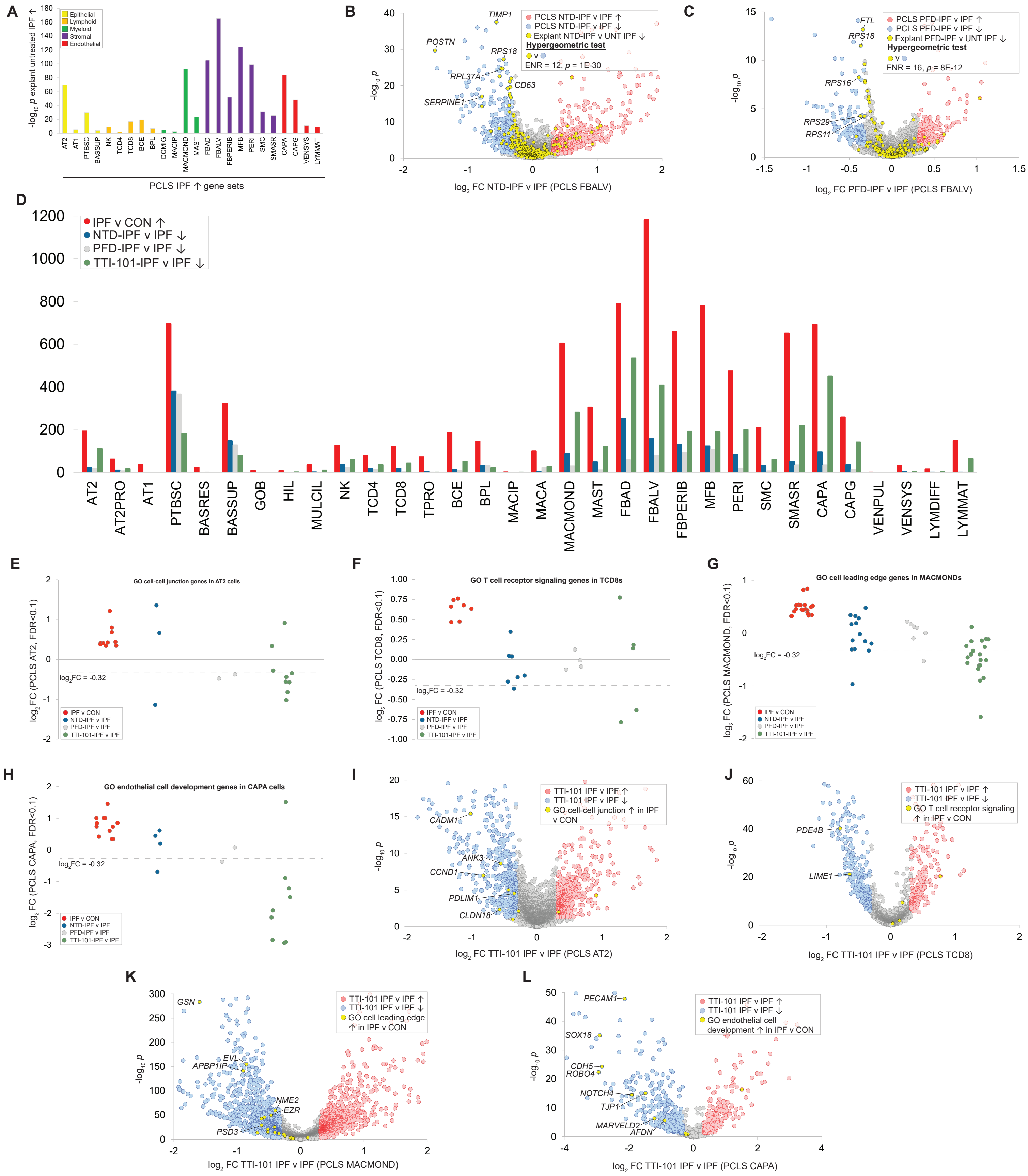

**Fig. S13.** **(A)** Bar chart showing enrichment -log₁₀*p* values for the intersection of PCLS and explant IPF-induced gene sets across common cell types. **(B)** Volcano plot of NTD-IPF vs. IPF-regulated genes in PCLS alveolar fibroblasts showing enrichment of explant NTD-repressed genes among PCLS NTD-repressed genes. **(C)** Volcano plot of PFD-IPF vs. IPF-regulated genes in PCLS alveolar fibroblasts showing enrichment of explant PFD-repressed genes among PCLS PFD-repressed genes. (**D)** Bar chart comparing nintedanib, pirfenidone and TTI-101 reversal of IPF-induced transcriptional programs across all PCLS cell types. Manhattan plots **E-H** compare numbers of IPF-induced and nintedanib-, pirfenidone- and TTI-101-repressed genes mapping to gap-prioritized GO terms from fig. S9: **(E)** cell-cell junction in AT2 cells; **(F)** T cell receptor signaling pathway in TCD8s; **(G)** cell leading edge in MACMONDs; **(H)** endothelial cell development in CAPAs. **I-L** represent TTI-101-IPF vs. IPF volcano plots showing TTI-101 repression of the indicated IPF-induced gap-prioritized GO term genes in: **(I)** cell-cell junction in AT2 cells; **(J)** T cell receptor signaling pathway in TCD8s; **(K)** cell leading edge in MACMONDs; and **(L)** endothelial cell development in CAPAs.

| Transcription factor classification | | | Explant FBALV | | | |
|---|---|---|---|---|---|---|
| | | | UNT IPF ← GRN $-\log_{10}p$ | $\log_2$FC (FDR<0.1) | | |
| Class | Family | Symbol | | IPF v CON | NTD v IPF | PFD v IPF |
| STAT domain | STAT factor | *STAT3* | | | | |
| BZIP factors | Jun factor | *JUN* | | | | |
| BZIP factors | Fos factor | *FOS* | | | | |
| BHLH factors | E2A-related factor | *TCF12* | | | | |
| Other transcription factors | Single GATA-type zinc-finger | *TRPS1* | | | | |
| BZIP factors | NF-E2-like factor | *NFE2L2* | | | | |
| BZIP factors | Jun factor | *JUNB* | | | | |
| C2H2 Zn finger factors | Kruppel-like | *KLF4* | | | | |
| BHLH factors | Ahr-like family | *AHR* | | | | |
| BHLH factors | Myc/Max factor | *MYC* | | | | |
| C2H2 Zn finger factors | ZNF99-like | *ZBTB16* | | | | |
| E2F/FOX | FOXP | *FOXP1* | | | | |
| C2H2 Zn finger factors | Kruppel-like | *KLF6* | | | | |
| BZIP factors | ATF-2-like factors | *CREB5* | | | | |
| E2F/FOX | FOXO | *FOXO1* | | | | |
| BHLH factors | Ahr-like family | *HIF1A* | | | | |
| Runt domain | Core-binding factor subunit | *RUNX1* | | | | |
| Runt domain | Core-binding factor subunit | *RUNX1T1* | | | | |
| Homeo domain | Zn finger E-box binding homeobox | *ZEB1* | | | | |
| Rel Homology Region | Early B-Cell Factor-related factor | *EBF1* | | | | |
| BHLH factors | Hairy-related | *BHLHE40* | | | | |
| BZIP factors | ATF-2-like factors | *ATF3* | | | | |
| BHLH factors | E2A-related factor | *TCF4* | | | | |

**Fig. S14.** Effect of nintedanib and pirfenidone on the 23 untreated IPF-induced explant alveolar fibroblast genes encoding IPF GRN TFs. Full data are in tables S5 (scRNA-Seq) and table S8 (GRN).
